# Cell Type-Specific Structural and Functional Signatures of APOE4, Age and Sex in the Mouse Anterior Olfactory Nucleus

**DOI:** 10.64898/2026.09.21.753183

**Authors:** Meigeng Hu, Alexander Kershaw, Sarah Brunson, Yaping Li, Dan Zhao, Shaolin Liu

## Abstract

As the strongest genetic risk factor for late-onset Alzheimer’s disease (AD), the human APOE ε4 allele alters neuronal physiology before overt pathology. However, how aging and sex modify these effects in the anterior olfactory nucleus (AON), an early AD-affected brain region, remains unclear. Here, we performed whole-cell recordings and post hoc morphological reconstructions of pyramidal cells (PCs) and interneurons (INs) in acute AON slices from adult and aged humanized APOE3 (E3) and APOE4 (E4) knock-in mice of both sexes with identification of context-dependent remodeling across structural, intrinsic, and synaptic domains. In PCs, morphology was broadly susceptible to APOE genotype, age, sex and their interactions. Functionally, aged E4 PCs exhibited a more depolarized resting membrane potential than aged E3 PCs. Additionally, afterhyperpolarization amplitude displayed a marked age × genotype interaction, reversing the direction of the E4–E3 difference from adulthood to aging. Excitatory synaptic event frequency showed a complex genotype × age × sex interaction, whereas inhibitory synaptic event amplitude was elevated in E4 mice. In contrast, INs displayed far more restricted alterations. Morphological changes were limited to dendritic length, which was modulated by age and sex specifically in E4 mice. Physiologically, genotype effects were confined to action potential amplitude, while aging primarily altered membrane properties and excitatory synaptic inputs. Together, these findings demonstrate that APOE genotype cell-specifically shapes AON neuronal properties through interactions with age and sex, suggesting that APOE4 establishes altered cellular states that influence how this circuit responds to subsequent disease-related stress.

**Significance Statement:** Carrying the APOE4 gene is the strongest genetic risk factor for late-onset Alzheimer’s disease (AD), and its effects on brain cells are not uniform, yet how they combine with aging and biological sex is largely unknown. We examined the anterior olfactory nucleus, a region affected very early in AD, and measured the morphology, physiological properties, and synaptic inputs of both excitatory and inhibitory neurons in mice carrying the human APOE3 or APOE4 gene. Rather than making neurons uniformly more or less active, APOE4 altered different properties in different cell types, and the direction of several effects depended on age and sex. Recognizing this context dependence may be necessary before cell-level changes can be linked to early disease risk.

## Introduction

Alzheimer’s disease (AD), the most common cause of dementia, is characterized by progressive memory loss and cognitive decline and imposes an immense burden worldwide(Lanctot et al., 2024). Despite extensive research, the cellular mechanisms underlying selective circuit vulnerability in AD remain incompletely understood.

Humans carry three common APOE alleles (ε2, ε3, ε4) encoding isoforms that differ by single amino acid substitutions. Among them, ε4 is the most potent genetic risk factor for sporadic, late-onset AD and acts with a strong gene dose dependency: heterozygous carriers show an approximately 2-3 fold elevation in risk, while homozygotes show a substantially greater increase in risk (12-15 fold) together with earlier symptom onset(Corder et al., 1993; Liu et al., 2013; Yamazaki et al., 2019).

Beyond its established effects on amyloid-β metabolism and tau-related pathology, ε4 has been linked to synaptic dysfunction, mitochondrial impairment, blood–brain barrier disruption, and altered neuroimmune responses(Bell et al., 2012; Shi et al., 2017; Pires and Rego, 2023), suggesting that it may alter brain homeostasis before overt neuropathology emerges. Consistent with this, ε4 has been associated with impaired synaptic plasticity, disrupted network oscillations, and altered excitability in cortical and limbic circuits(Bu, 2009; Huang and Mahley, 2014). Humanized APOE knock-in mice express human ε3 or ε4 in place of the mouse gene without developing the plaques and tangles seen in AD brains, allowing APOE-dependent effects to be examined separately from established amyloid and tau pathology(Sullivan et al., 1997; Sullivan et al., 2004; Wang et al., 2005; Nuriel et al., 2017; Onos et al., 2024).

APOE genotype does not act on a static background. Age is the strongest non-genetic risk factor for AD and is accompanied by synaptic loss, metabolic decline, and neuroinflammatory priming(Mattson and Arumugam, 2018). Sex differences are increasingly recognized: women, particularly ε4 carriers, have shown greater AD risk or faster cognitive decline in several cohorts(Farrer et al., 1997; Holland et al., 2013; Altmann et al., 2014; Riedel et al., 2016; Neu et al., 2017). These three factors converge on the same neurons, yet are usually studied one or two at a time, leaving the cellular substrates of their interaction poorly defined.

The anterior olfactory nucleus (AON) is an evolutionarily conserved cortical relay between the olfactory bulb and higher-order limbic structures(Haberly and Price, 1977; Brunert et al., 2023) and communicates extensively with cortical regions involved in memory and behavioral regulation(Brunjes et al., 2005; Aqrabawi and Kim, 2018; Zhou et al., 2019; Bhattarai et al., 2022). Despite evidence of early tau pathology and neuronal degeneration in the AON(Franks et al., 2015; Murray et al., 2020), what is known about the functional impact of ε4 in this region comes principally from our recent study showing APOE genotype-, age-and sex-associated alterations in neuronal and network activity with high-density extracellular recording in awake mice(Uzun et al., 2025). Specifically, adult ε4 mice had lower spontaneous spike output than ε3 animals regardless of sex, and adult females had higher spike output than adult males across genotypes. Those recordings establish that all three AD risk factors (APOE4, aging, and sex) shape AON output, but the cellular and synaptic mechanisms remain elusive.

Here we addressed this question using whole-cell recordings combined with post hoc morphological reconstruction of biocytin-filled neurons in acute brain slices from adult and aged humanized APOE knock-in mice homozygous for ε3 or ε4 of both sexes. We quantified soma area and dendritic architecture, resting membrane potential (RMP), input resistance (R_in_), action potential (AP) and afterhyperpolarization (AHP) amplitude, and spontaneous excitatory (sEPSCs) and inhibitory postsynaptic currents (sIPSCs) to determine whether APOE genotype, age, and sex converge on common mechanisms or remodel distinct properties in a cell type–specific manner. Our findings support the latter: APOE genotype affected distinct cellular parameters across neuron types, with several effects contingent on age and sex and, in some cases, differing in direction between sexes. Thus, APOE4 produces context-dependent remodeling of AON neurons rather than a uniform shift in excitability.

## Materials and Methods

### Animals

The breeder pairs for humanized ε3 (B6.Cg-APOEem2(APOE*)Adiuj/J, strain# 029018) and ε4 knock-in mice (B6(SJL)-APOEtm1.1(APOE*4)Adiuj/J, strain# 027894) were purchased from the Jackson Laboratory (Bar Harbor, ME, USA) and bred in-house at the University of Georgia. Animals were housed in a specific pathogen-free facility under a 12-hour light/dark cycle with ad libitum access to chow and water. Only homozygous ε4+/+ (E4) and ε3+/+ (E3) mice of both sexes at two ages (adult: 27 weeks; aged: 76 weeks) were used for experiments. Data were collected from animals in eight groups stratified by age, sex, and strain: (1) aged E4 females (n=7 mice); (2) aged E4 males (n=9 mice); (3) adult E4 females (n=8 mice); (4) adult E4 males (n=7 mice); (5) aged E3 females (n=6 mice); (6) aged E3 males (n=8 mice); (7) adult E3 females (n=9 mice); (8) adult E3 males (n=6 mice). All procedures were conducted in accordance with National Institutes of Health (NIH) guidelines for the care and use of laboratory animals and were approved by the Institutional Animal Care and Use Committee of the University of Georgia.

### Slice Preparation

Acute brain slices (300 *μ*m thick) containing the AON were prepared from E3 and E4 mice as previously described(Liu and Liu, 2018; Liu, 2020). Briefly, animals were deeply anesthetized with isoflurane before decapitation. Coronal sections were cut using a VT1200S vibratome (Leica Microsystems, Germany) in ice-cold, oxygenated (95% O₂ / 5% CO₂) NMDG-based artificial cerebrospinal fluid (NMDG-aCSF) containing (in mM): 90 NMDG, 2.5 KCl, 1.2 NaH_2_PO_4_, 30 NaHCO_3_, 25 glucose, 20 NaHEPES, 5 Sodium ascorbate, 2 Thiourea, 3 Sodium pyruvate, 10 MgSO_4_, 0.5 CaCl_2_ (pH 7.40, 300 mOsm). After 30 min of incubation in NMDG aCSF at 30°C, slices were then transferred to HEPES-based aCSF at room temperature (RT) until they were used for recordings.

HEPES-based aCSF was continuously bubbled with 95% O_2_–5% CO_2_ and had the composition (in mM): 92 NaCl, 2.5 KCl, 1.2 NaH_2_PO_4_, 30 NaHCO_3_, 25 glucose, 20 NaHEPES, 2 Sodium ascorbate, 2 Thiourea, 3 Sodium pyruvate, 2 MgSO_4_, 2 CaCl_2_ (pH 7.40, 300 mOsm). During the experiments, slices were perfused at 2.5 ml/min with recording aCSF, which was equilibrated with 95% O_2_–5% CO_2_ and warmed to 30°C and had the composition (in mM): 125 NaCl, 2.5 KCl, 1.25 NaH_2_PO_4_, 25 NaHCO_3_, 10 glucose, 2 MgSO_4_, 2 CaCl_2_ (pH 7.40, 300 mOsm).

### Electrophysiology

Whole-cell patch-clamp recordings were performed from neurons in the AON visualized with an Eclipse FN1 fixed-stage upright microscope (Nikon, Japan) equipped with near-infrared differential interference contrast (IR-DIC) optics. The AON is anatomically divided into two subregions: pars externa and pars principalis. The pars externa situates between the olfactory bulb and the pars principalis to mainly serve as an inter-bulb connection hub. On the other hand, the pars principalis as the largest cellular body of AON has four subdivisions: lateral, medial, dorsal, and ventroposterior (Brunjes et al., 2005). Due the cytoarchitectural and potentially functional homogeneity in the dorsolateral regions (Meyer et al., 2006), we purposely confined our recordings to these two subregions to minimize regional variations. Neurons were preselected for recording based on soma location and dendritic projection patterns(Brunjes and Kenerson, 2010; Kay and Brunjes, 2014), with cell identity subsequently verified by electrophysiological signatures and post hoc morphological reconstruction.

To enable post hoc visualization of somatodendritic architecture, biocytin (0.2%) was included in the internal solution. Signals were acquired in either current-or voltage-clamp mode using the Sutter IPA (Integrated Patch Amplifier) and SutterPatch® Data Acquisition Software (v3.1). Data were sampled and filtered at 5 kHz using the IPA’s built-in filter, with automated series resistance and electrode compensation applied during acquisition.

Patch pipettes (5–7 MΩ) were pulled from thin-walled glass capillaries with filament (Sutter Instrument, Novato, CA). The pipette internal solution contained (in mM): 117 K-gluconate, 10 KCl, 10 HEPES, 4 EGTA, 12 KOH, 0.5 CaCl₂, 3 Mg-ATP, 0.3 Na₂-GTP, and 7 Na₂-phosphocreatine (pH adjusted to 7.26, 290 mOsm). Data analysis was conducted in SutterPatch as described below.

### Histology and Confocal Imaging

Following electrophysiological recordings, brain slices containing biocytin (0.2%, w/v)-filled neurons were transferred immediately to 4% paraformaldehyde (PFA) and fixed at RT for 30 minutes. Slices were then rinsed three times (3 min/each) in 0.05 M phosphate-buffered saline (PBS) and incubated for 1 hour in blocking solution (10% normal donkey serum, 2% bovine serum albumin, and 0.25% Triton X-100 in 0.05 M PBS) on a shaker.

Subsequently, slices were incubated for 5 hours at RT in the dark on a shaker with fresh blocking solution containing streptavidin-conjugated Cy3 (1.8 *μ*g/ml). After staining, slices were rinsed three times (5 min/each) in PBS and then incubated with 0.05 M PBS containing 4′,6-diamidino-2-phenylindole (DAPI; 50 *μ*g/ml) for 10 minutes at RT in the dark to label nuclei. Slices were rewashed three times (5 min/each), then wet-mounted and coverslipped with mounting medium.

Confocal image stacks were acquired using a Zeiss LSM900 microscope equipped with a 40× oil-immersion objective. Z-stacks were collected with a step size of 1 *μ*m and an XY resolution of 0.21 *μ*m/pixel. Morphological reconstruction and quantification were performed on z-projected images using Fiji as described below.

### Morphological Characterization

Morphological analysis was performed on biocytin-filled neurons in the AON-containing brain slices visualized with streptavidin-Cy3-staining following whole-cell patch-clamp recordings. Maximum-intensity projections were generated in FIJI (v2.16.0). Soma size was quantified on maximum-intensity projections by outlining the cell body with the freehand selection tool in FIJI and measuring the enclosed two-dimensional projected area, which may not fully capture three-dimensional changes in soma volume (Kay and Brunjes, 2014).

For dendritic reconstruction, images were converted to 8-bit and traced manually using the Simple Neurite Tracer (SNT) plugin in FIJI in the XY view. Primary dendrites were defined as processes emerging directly from the soma, and branches as processes arising from a primary dendrite or from a higher-order branch; branches of all orders were pooled without further stratification. Path lengths were obtained from the SNT Path Manager (Tag > Morphometry > Lengths). Because tracing was performed in two dimensions, dendritic lengths represent projected path lengths. Total dendritic length (TDL) was calculated as the summed length of all primary dendrites and branches; branch number as the number of branch segments, excluding primary dendrites; and branch length as the mean length of individual branch segments, excluding primary dendrites.

## Data Analysis

Electrophysiological data were analyzed using Igor Pro 9.05 (WaveMetrics). R_in_ was measured immediately upon achieving whole-cell configuration in voltage clamp with a 40 ms hyperpolarizing 5 mV voltage step whereas RMP was determined in current clamp without current injection.

To assess single action potential (AP) characteristics including amplitude and afterhyperpolarization (AHP), current injections were delivered in 30 pA (300 ms) increments to evoke neuronal firing. AP amplitude was measured as the voltage difference between the spike threshold and its peak. AHP was measured as the voltage difference between the AP threshold and the most negative point of the subsequent hyperpolarization. sEPSC and sIPSC detection was performed in Igor Pro 9.05 using a semi-automated threshold-based algorithm followed by manual confirmation. Amplitude was defined as the peak current relative to baseline. Instantaneous frequencies were calculated as the inverse of the interval between any two consecutive events, as determined by the detection algorithm. Cumulative probability distributions of sEPSC and sIPSC amplitudes and frequencies were generated using a custom-written algorithm in Microsoft Excel.

### Experimental Design and Statistical Analyses

The number of animals per group is provided in Animals under Methods section; the number of cells analyzed per group is shown in the figure legends. Group data are presented as mean ± SEM. Each parameter was analyzed by three-way ANOVA with APOE genotype, age, and sex as between-subject factors. Post hoc pairwise comparisons with Holm–Bonferroni correction were performed only when a main effect or interaction reached statistical significance. The relationship between input resistance and dendritic branch number was assessed by Pearson correlation. Statistical significance was set at p < 0.05. Analyses were performed in Origin 2025 (OriginLab).

## Results

### Effects of APOE genotype, age, and sex on neuronal morphology

Multiple lines of evidence have suggested that neuronal morphology is sculpted by APOE genotypes and age. For example, E4 KI mice showed smaller pyramidal cells (PCs) in the hippocampal CA3 region than E3 KI mice at young ages, an atrophy that was also observed in E3 neurons with aging(Tabuena et al., 2026). Additionally, embryonic hippocampal neurons exposed to ApoE4 astrocyte-conditioned medium showed shorter neurites than those exposed to ApoE3-conditioned medium(Thorwald et al., 2025). E4 has been associated with progressive loss of hilar GABAergic interneurons and age-related inhibitory dysfunction in the hippocampus(Li et al., 2009; Tabuena et al., 2026). To assess whether AON neurons are subject to these types of modulation, we quantified and compared the two-dimensional projected area of the soma (soma area) alongside dendritic metrics, including the number of branches, average branch length, and total dendritic length (TDL), for both pyramidal cells (PCs) and interneurons (INs) among groups of animals stratified by APOE genotype, age and sex. Because the AON contains morphologically and electrophysiologically distinct pyramidal-cell and interneuron classes(Brunjes and Kenerson, 2010; Kay and Brunjes, 2014), PCs and INs were analyzed separately.

### Soma size

All recorded cells were reconstructed by staining the pipette-filled biocytin with CY3-conjugated streptavidin to verify their identity (Fig 1A & 1B). Three-way ANOVA revealed strong influence of age on PCs (F(1,126) = 12.79, p = 0.00049; Tab. 1), with aged E4 mice exhibiting smaller somata than their adult E4 counterparts of both sexes (males: t = −3.77, p = 0.00025; females: t = −3.26, p = 0.00146; Fig. 1C). These age-related differences were absent in E3 mice, consistent with a significant age × genotype interaction (F(1,126) = 11.42, p = 0.00097; Tab. 1). This interaction was further reflected by the opposite direction of the genotype effect depending on age and sex: aged E4 males had smaller somata than aged E3 males (t = −2.57, p = 0.01121), whereas adult E4 females had larger somata than adult E3 females (t = 2.29, p = 0.02363; Fig. 1C). In contrast, no differences were observed in INs (Fig 1D; Tab. 1). These data demonstrate that soma area in PCs depended on an age × genotype interaction: soma area was larger in adult E4 than adult E3 females, whereas in aged animals it was smaller in E4 than E3 males. Soma area was also smaller in aged than adult E4 mice of both sexes.

**Figure 1.**
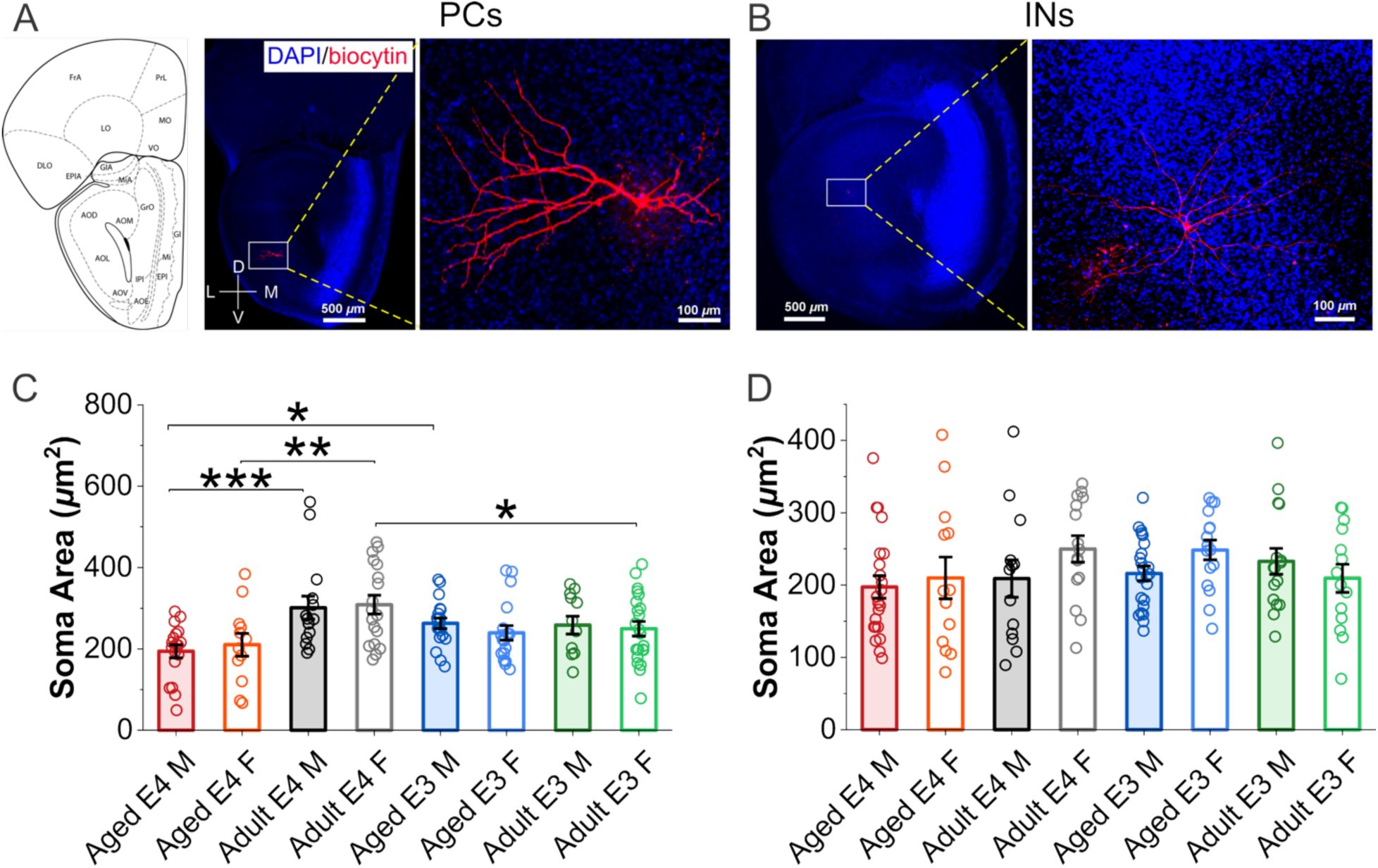
Effects of genotype, age, and sex on the soma size. **A & B**, confocal images of coronal AON sections from an aged male (A) or adult female (B) APOE4 mouse. Left: biocytin-filled pyramidal cell (PC, A) or interneuron (IN, B) were visualized using streptavidin-CY3 (red) with DAPI counterstaining (blue). Right: zoomed-in image of the white-framed area from the left to highlight the difference in cell morphology. **C&D**, comparison of soma sizes (measured as two-dimensional projected area) of PCs (C) and INs (D) across groups stratified by genotype, age, and sex. Sample sizes (cells/group): aged APOE4 (E4) males (n=18 PCs & 22 INs), aged E4 females (n=12 PCs & 13 INs), adult E4 males (n=15 PCs & 13 INs), adult E4 females (n=18 PCs & 14 INs), aged APOE3 (E3) males (n=19 PCs & 23 INs), aged E3 females (n=18 PCs & 16 INs), adult E3 males (n=12 PCs & 16 INs), and adult E3 females (n=22 PCs & 14 INs). Bar graphs represent group means ± SEM with individual data points represented by symbols. Three-way ANOVA with Holm–Bonferroni correction, *p < 0.05, **p < 0.01, ***p < 0.001, ****p < 0.0001.

**Table 1.** Three-way ANOVA of the effects of APOE genotype, age, and sex on morphological, intrinsic, and synaptic properties of PCs and INs in the AON, with post hoc pairwise comparisons.

| Figure | panel | Factor | Statistic | p value | post-hoc comparison, t value, and p value |
| --- | --- | --- | --- | --- | --- |
| Figure 1 | C | Age<br>Sex<br>Genotype<br>Age x Sex<br>Age x Genotype<br>Sex x Genotype<br>Age x Sex x Genotype | F (1, 126) = 12.793<br>F (1, 126) = 0.022<br>F (1, 126) = 0.004<br>F (1, 126) = 0.01<br>F (1, 126) = 11.424<br>F (1, 126) = 0.898<br>F (1, 126) = 0.152 | p = 0.00049<br>p = 0.88265<br>p = 0.94915<br>p = 0.92031<br>p = 0.00097<br>p = 0.34518<br>p = 0.69746 | E4 Aged M vs. E3 Aged M: t = -2.574, p = 0.01121<br>E4 Aged M vs. E4 Adult M: t = -3.771, p = 0.00025<br>E4 Aged F vs. E4 Adult F: t = -3.255, p = 0.00146<br>E4 Adult F vs. E3 Adult F: t = 2.291, p = 0.02363 |
|  | D | Age<br>Sex<br>Genotype<br>Age x Sex<br>Age x Genotype<br>Sex x Genotype<br>Age x Sex x Genotype | F (1, 123) = 0.379<br>F (1, 123) = 1.549<br>F (1, 123) = 0.526<br>F (1, 123) = 0.225<br>F (1, 123) = 2.165<br>F (1, 123) = 0.813<br>F (1, 123) = 2.768 | p = 0.53936<br>p = 0.21571<br>p = 0.46957<br>p = 0.63641<br>p = 0.14374<br>p = 0.36911<br>p = 0.09873 | N/A |
| Figure 2 | A | Age<br>Sex<br>Genotype<br>Age x Sex<br>Age x Genotype<br>Sex x Genotype<br>Age x Sex x Genotype | F (1, 126) = 7.704<br>F (1, 126) = 11.5<br>F (1, 126) = 2.733<br>F (1, 126) = 11.045<br>F (1, 126) = 0.016<br>F (1, 126) = 7.848<br>F (1, 126) = 2.247 | p = 0.00635<br>p = 0.00093<br>p = 0.10078<br>p = 0.00116<br>p = 0.89846<br>p = 0.00589<br>p = 0.13636 | E4 Aged M vs. E4 Aged F: t = 3.914, p = 0.00015<br>E3 Aged M vs. E3 Aged F: t = 2.994, p = 0.00332<br>E4 Aged F vs. E4 Adult F: t = -2.308, p = 0.02261<br>E3 Aged F vs. E3 Adult F: t = -4.277, p < 0.0001<br>E4 Adult M vs. E4 Adult F: t = 2.273, p = 0.02469<br>E3 Adult M vs. E3 Adult F: t = -2.146, p = 0.03379<br>E4 Adult F vs. E3 Adult F: t = -3.335, p = 0.00112 |
|  | B | Age<br>Sex<br>Genotype<br>Age x Sex<br>Age x Genotype<br>Sex x Genotype<br>Age x Sex x Genotype | F (1, 123) = 0.738<br>F (1, 123) = 0.071<br>F (1, 123) = 1.085<br>F (1, 123) = 0.481<br>F (1, 123) = 0.66<br>F (1, 123) = 0.703<br>F (1, 123) = 2.797 | p = 0.39199<br>p = 0.79033<br>p = 0.29958<br>p = 0.48931<br>p = 0.41799<br>p = 0.40342<br>p = 0.097 | N/A |
|  | C | Age<br>Sex<br>Genotype<br>Age x Sex<br>Age x Genotype<br>Sex x Genotype<br>Age x Sex x Genotype | F (1, 126) = 10.293<br>F (1, 126) = 8.858<br>F (1, 126) = 0.129<br>F (1, 126) = 5.028<br>F (1, 126) = 0.127<br>F (1, 126) = 6.888<br>F (1, 126) = 0.313 | p = 0.00169<br>p = 0.0035<br>p = 0.71974<br>p = 0.0267<br>p = 0.72269<br>p = 0.00975<br>p = 0.57656 | E4 Aged M vs. E4 Aged F: t = 3.556, p = 0.00053<br>E4 Aged F vs. E4 Adult F: t = -2.214, p = 0.02862<br>E3 Aged F vs. E3 Adult F: t = -3.645, p = 0.00039<br>E4 Adult M vs. E4 Adult F: t = 2.039, p = 0.04351 |
|  | D | Age<br>Sex<br>Genotype<br>Age x Sex<br>Age x Genotype<br>Sex x Genotype<br>Age x Sex x Genotype | F (1, 123) = 5.901<br>F (1, 123) = 3.97<br>F (1, 123) = 0.045<br>F (1, 123) = 0.104<br>F (1, 123) = 1.031<br>F (1, 123) = 0.0<br>F (1, 123) = 2.147 | p = 0.01658<br>p = 0.04853<br>p = 0.83195<br>p = 0.74758<br>p = 0.31195<br>p = 0.99986<br>p = 0.14544 | E4 Aged F vs. E4 Adult F: t = -2.504, p = 0.01357 |
|  | E | Age<br>Sex<br>Genotype<br>Age x Sex<br>Age x Genotype<br>Sex x Genotype<br>Age x Sex x Genotype | F (1, 126) = 0.103<br>F (1, 126) = 0.672<br>F (1, 126) = 7.037<br>F (1, 126) = 1.421<br>F (1, 126) = 3.079<br>F (1, 126) = 0.623<br>F (1, 126) = 0.487 | p = 0.74903<br>p = 0.41377<br>p = 0.00901<br>p = 0.23546<br>p = 0.08173<br>p = 0.43142<br>p = 0.48635 | E4 Aged M vs. E3 Aged M: t = 2.489, p = 0.01413<br>E4 Aged F vs. E3 Aged F: t = 2.107, p = 0.03706 |
|  | F | Age<br>Sex<br>Genotype<br>Age x Sex<br>Age x Genotype<br>Sex x Genotype<br>Age x Sex x Genotype | F (1, 123) = 4.436<br>F (1, 123) = 4.165<br>F (1, 123) = 2.797<br>F (1, 123) = 1.842<br>F (1, 123) = 1.233<br>F (1, 123) = 0.617<br>F (1, 123) = 0.01 | p = 0.03722<br>p = 0.04341<br>p = 0.09696<br>p = 0.17716<br>p = 0.26895<br>p = 0.43381<br>p = 0.9195 | E4 Aged M vs. E4 Aged F: t = -2.258, p = 0.0257<br>E4 Aged M vs. E4 Adult M: t = -2.464, p = 0.01512 |
| Figure 3 | A | Age<br>Sex<br>Genotype<br>Age x Sex<br>Age x Genotype<br>Sex x Genotype<br>Age x Sex x Genotype | F (1, 126) = 1.25<br>F (1, 126) = 0.048<br>F (1, 126) = 6.098<br>F (1, 126) = 0.847<br>F (1, 126) = 9.865<br>F (1, 126) = 0.26<br>F (1, 126) = 1.12 | p = 0.26561<br>p = 0.82647<br>p = 0.01487<br>p = 0.35919<br>p = 0.0021<br>p = 0.61108<br>p = 0.29188 | E4 Aged M vs. E3 Aged M: t = 3.407, p = 0.00088<br>E4 Aged M vs. E4 Adult M: t = 3.247, p = 0.0015<br>E4 Aged F vs. E3 Aged F: t = 2.471, p = 0.0148 |
|  | B | Age<br>Sex<br>Genotype<br>Age x Sex<br>Age x Genotype<br>Sex x Genotype<br>Age x Sex x Genotype | F (1, 123) = 5.124<br>F (1, 123) = 0.692<br>F (1, 123) = 0.025<br>F (1, 123) = 2.915<br>F (1, 123) = 3.03<br>F (1, 123) = 3.394<br>F (1, 123) = 1.932 | p = 0.02535<br>p = 0.40699<br>p = 0.87571<br>p = 0.09029<br>p = 0.08424<br>p = 0.06783<br>p = 0.167 | E3 Aged F vs. E3 Adult F: t = -3.578, p = 0.0005 |
|  | C | Age<br>Sex<br>Genotype<br>Age x Sex<br>Age x Genotype<br>Sex x Genotype<br>Age x Sex x Genotype | F (1, 126) = 0.836<br>F (1, 126) = 0.0<br>F (1, 126) = 2.497<br>F (1, 126) = 0.059<br>F (1, 126) = 4.727<br>F (1, 126) = 0.029<br>F (1, 126) = 3.837 | p = 0.36228<br>p = 0.99153<br>p = 0.11656<br>p = 0.80914<br>p = 0.03156<br>p = 0.86496<br>p = 0.05236 | E4 Aged F vs. E4 Adult F: t = -2.346, p = 0.02053<br>E3 Aged F vs. E3 Adult F: t = 1.981, p = 0.04975<br>E4 Adult F vs. E3 Adult F: t = 3.173, p = 0.00189 |
|  | D | Age<br>Sex<br>Genotype<br>Age x Sex<br>Age x Genotype<br>Sex x Genotype<br>Age x Sex x Genotype | F (1, 123) = 0.139<br>F (1, 123) = 4.645<br>F (1, 123) = 0.764<br>F (1, 123) = 0.079<br>F (1, 123) = 0.934<br>F (1, 123) = 0.318<br>F (1, 123) = 0.8 | p = 0.71002<br>p = 0.0331<br>p = 0.3837<br>p = 0.77979<br>p = 0.33584<br>p = 0.57376<br>p = 0.37277 | E4 Aged M vs. E4 Aged F: t = 2.052, p = 0.04227 |
| Figure 4 | C | Age<br>Sex<br>Genotype<br>Age x Sex<br>Age x Genotype<br>Sex x Genotype<br>Age x Sex x Genotype | F (1, 126) = 7.4<br>F (1, 126) = 4.351<br>F (1, 126) = 2.98<br>F (1, 126) = 0.446<br>F (1, 126) = 0.64<br>F (1, 126) = 0.37<br>F (1, 126) = 0.133 | p = 0.00744<br>p = 0.03901<br>p = 0.08672<br>p = 0.50525<br>p = 0.42534<br>p = 0.54332<br>p = 0.71606 | N/A |
|  | D | Age<br>Sex<br>Genotype<br>Age x Sex<br>Age x Genotype<br>Sex x Genotype<br>Age x Sex x Genotype | F (1, 123) = 3.268<br>F (1, 123) = 0.512<br>F (1, 123) = 2.179<br>F (1, 123) = 1.097<br>F (1, 123) = 0.069<br>F (1, 123) = 4.403<br>F (1, 123) = 0.105 | p = 0.07307<br>p = 0.47545<br>p = 0.14249<br>p = 0.29692<br>p = 0.79372<br>p = 0.03791<br>p = 0.74664 | E4 Aged M vs. E3 Aged M: t = -2.572, p = 0.0113<br>E4 Aged M vs. E4 Aged F: t = -2.206, p = 0.02925 |
|  | E | Age<br>Sex<br>Genotype<br>Age x Sex<br>Age x Genotype<br>Sex x Genotype<br>Age x Sex x Genotype | F (1, 126) = 3.786<br>F (1, 126) = 0.434<br>F (1, 126) = 0.447<br>F (1, 126) = 1.071<br>F (1, 126) = 29.322<br>F (1, 126) = 0.0<br>F (1, 126) = 0.401 | p = 0.05392<br>p = 0.51124<br>p = 0.50506<br>p = 0.30269<br>p < 0.0001<br>p = 0.98282<br>p = 0.5279 | E4 Aged M vs. E3 Aged M: t = -2.964, p = 0.00363<br>E4 Aged M vs. E4 Adult M: t = -4.699, p < 0.0001<br>E4 Aged F vs. E3 Aged F: t = -2.019, p = 0.04562<br>E4 Aged F vs. E4 Adult F: t = -2.779, p = 0.00628<br>E3 Aged F vs. E3 Adult F: t = 2.216, p = 0.02847<br>E4 Adult M vs. E3 Adult M: t = 3.166, p = 0.00194<br>E4 Adult F vs. E3 Adult F: t = 3.108, p = 0.00233 |
|  | F | Age<br>Sex<br>Genotype<br>Age x Sex<br>Age x Genotype<br>Sex x Genotype<br>Age x Sex x Genotype | F (1, 123) = 5.43<br>F (1, 123) = 0.532<br>F (1, 123) = 0.925<br>F (1, 123) = 0.011<br>F (1, 123) = 2.398<br>F (1, 123) = 0.024<br>F (1, 123) = 0.007 | p = 0.02143<br>p = 0.46725<br>p = 0.33806<br>p = 0.91478<br>p = 0.1241<br>p = 0.87591<br>p = 0.93255 | E4 Aged M vs. E4 Adult M: t = -2.145, p = 0.03389 |
| Figure 5 | F | Age<br>Sex<br>Genotype<br>Age x Sex<br>Age x Genotype<br>Sex x Genotype<br>Age x Sex x Genotype | F (1, 103) = 1.71<br>F (1, 103) = 1.948<br>F (1, 103) = 5.038<br>F (1, 103) = 5.518<br>F (1, 103) = 3.8<br>F (1, 103) = 4.868<br>F (1, 103) = 6.075 | p = 0.1939<br>p = 0.16586<br>p = 0.02694<br>p = 0.02074<br>p = 0.05396<br>p = 0.02959<br>p = 0.01537 | E4 Aged M vs. E3 Aged M: t = -2.41, p = 0.01775<br>E4 Aged M vs. E4 Aged F: t = -1.994, p = 0.04882<br>E4 Aged M vs. E4 Adult M: t = -2.871, p = 0.00497<br>E4 Aged F vs. E3 Aged F: t = -1.995, p = 0.04872<br>E4 Aged F vs. E4 Adult F: t = 2.038, p = 0.04416<br>E4 Adult M vs. E3 Adult M: t = 2.198, p = 0.03019<br>E4 Adult M vs. E4 Adult F: t = 2.896, p = 0.00462<br>E4 Adult F vs. E3 Adult F: t = -2.626, p = 0.00995 |
|  | G | Age<br>Sex<br>Genotype<br>Age x Sex<br>Age x Genotype<br>Sex x Genotype<br>Age x Sex x Genotype | F (1, 120) = 8.782<br>F (1, 120) = 0.141<br>F (1, 120) = 2.732<br>F (1, 120) = 1.188<br>F (1, 120) = 0.165<br>F (1, 120) = 1.063<br>F (1, 120) = 0.885 | p = 0.00367<br>p = 0.70796<br>p = 0.10099<br>p = 0.27796<br>p = 0.68522<br>p = 0.30453<br>p = 0.34867 | E3 Aged M vs. E3 Adult M: t = -3.021, p = 0.00308 |
| Figure 6 | E | Age<br>Sex<br>Genotype<br>Age x Sex<br>Age x Genotype<br>Sex x Genotype<br>Age x Sex x Genotype | F (1, 103) = 6.547<br>F (1, 103) = 2.372<br>F (1, 103) = 0.558<br>F (1, 103) = 0.001<br>F (1, 103) = 0.167<br>F (1, 103) = 0.098<br>F (1, 103) = 2.465 | p = 0.01196<br>p = 0.12661<br>p = 0.45686<br>p = 0.97768<br>p = 0.68375<br>p = 0.75488<br>p = 0.1195 | E3 Aged F vs. E3 Adult F: t = -2.522, p = 0.0132 |
|  | F | Age<br>Sex<br>Genotype<br>Age x Sex<br>Age x Genotype<br>Sex x Genotype<br>Age x Sex x Genotype | F (1, 120) = 13.181<br>F (1, 120) = 0.464<br>F (1, 120) = 6.082<br>F (1, 120) = 0.916<br>F (1, 120) = 0.014<br>F (1, 120) = 0.866<br>F (1, 120) = 0.454 | p = 0.00042<br>p = 0.49698<br>p = 0.01507<br>p = 0.34043<br>p = 0.90519<br>p = 0.35393<br>p = 0.50161 | E4 Aged F vs. E4 Adult F: t = -2.478, p = 0.01458<br>E3 Aged F vs. E3 Adult F: t = -2.046, p = 0.04296 |
| Figure 7 | D | Age<br>Sex<br>Genotype<br>Age x Sex<br>Age x Genotype<br>Sex x Genotype<br>Age x Sex x Genotype | F (1, 119) = 0.012<br>F (1, 119) = 0.007<br>F (1, 119) = 0.0<br>F (1, 119) = 0.1<br>F (1, 119) = 0.38<br>F (1, 119) = 1.195<br>F (1, 119) = 0.647 | p = 0.9142<br>p = 0.93248<br>p = 0.98797<br>p = 0.7522<br>p = 0.53881<br>p = 0.27648<br>p = 0.42296 | N/A |
|  | E | Age<br>Sex<br>Genotype<br>Age x Sex<br>Age x Genotype<br>Sex x Genotype<br>Age x Sex x Genotype | F (1, 105) = 2.203<br>F (1, 105) = 0.665<br>F (1, 105) = 0.013<br>F (1, 105) = 0.384<br>F (1, 105) = 0.859<br>F (1, 105) = 0.922<br>F (1, 105) = 3.614 | p = 0.14073<br>p = 0.41672<br>p = 0.90907<br>p = 0.53672<br>p = 0.35609<br>p = 0.33927<br>p = 0.06002 | N/A |
| Figure 8 | C | Age<br>Sex<br>Genotype<br>Age x Sex<br>Age x Genotype<br>Sex x Genotype<br>Age x Sex x Genotype | F (1, 119) = 0.127<br>F (1, 119) = 2.353<br>F (1, 119) = 14.058<br>F (1, 119) = 1.035<br>F (1, 119) = 1.799<br>F (1, 119) = 0.013<br>F (1, 119) = 0.272 | p = 0.72237<br>p = 0.12767<br>p = 0.00028<br>p = 0.31101<br>p = 0.18245<br>p = 0.91031<br>p = 0.60308 | E4 Aged M vs. E3 Aged M: t = 3.091, p = 0.00249<br>E4 Aged F vs. E3 Aged F: t = 2.292, p = 0.02366 |
|  | D | Age<br>Sex<br>Genotype<br>Age x Sex<br>Age x Genotype<br>Sex x Genotype<br>Age x Sex x Genotype | F (1, 105) = 0.436<br>F (1, 105) = 0.178<br>F (1, 105) = 5.013<br>F (1, 105) = 0.126<br>F (1, 105) = 0.254<br>F (1, 105) = 0.127<br>F (1, 105) = 0.368 | p = 0.51072<br>p = 0.67423<br>p = 0.02727<br>p = 0.72383<br>p = 0.61541<br>p = 0.72228<br>p = 0.54558 | N/A |

### Dendritic architecture

Given that somatic size was reshaped by the interaction of genotype and age, we next asked whether dendritic arbors arising from these somas were similarly affected. Thus, number of branches, TDL, and average branch length in the same sets of PCs and INs were quantified. Consistent with data on soma size, number of branches was shaped by age (F(1,126) = 7.70, p = 0.00635) and sex (F(1,126) = 11.50, p = 0.00093), with significant age × sex (F(1,126) = 11.04, p = 0.00116) and genotype × sex (F(1,126) = 7.85, p = 0.00589) interactions in PCs (Fig 2A) but not in INs (Fig 2B). Aging reduced number of branches in PCs of both E3 (from 19.9 ± 1.3 in adult, n = 22 to 11.8 ± 1.5 in aged, n = 18, a 40.6% reduction; t = −4.277, p < 0.0001) and E4 females (from 13.6 ± 1.3 in adult, n = 18 to 8.5 ± 1.5 in aged, n = 12, a 37.5% reduction; t = −2.308, p = 0.02261) but not males. In contrast, sex effects were broadly present. Specifically, males possessed more branches than females in the aged E4 (17.2 ± 1.7, n = 18 vs 8.5 ± 1.5, n = 12; t = 3.91, p = 0.00015), aged E3 (17.7 ± 1.4, n = 19 vs 11.8 ± 1.5, n = 18; t = 2.99, p = 0.00332), and adult E4 (18.3 ± 1.7, n = 15 vs 13.6 ± 1.3, n = 18; t = 2.27, p = 0.02469) groups, whereas this sex difference reversed in adult E3 animals, in which females exhibit more branches than males (19.9 ± 1.3, n = 22 vs 15.3 ± 1.5, n = 12; t = −2.15, p = 0.03379; Fig. 2A). In line with the genotype × sex interaction, adult E4 females possessed fewer branches than adult E3 females (13.6 ± 1.3 vs 19.9 ± 1.3; t =−3.34, p = 0.00112; Fig. 2A). In PCs, branch number was therefore shaped by age and sex and by the interaction between sex and genotype, with the genotype difference present in adult females.

**Figure 2.**
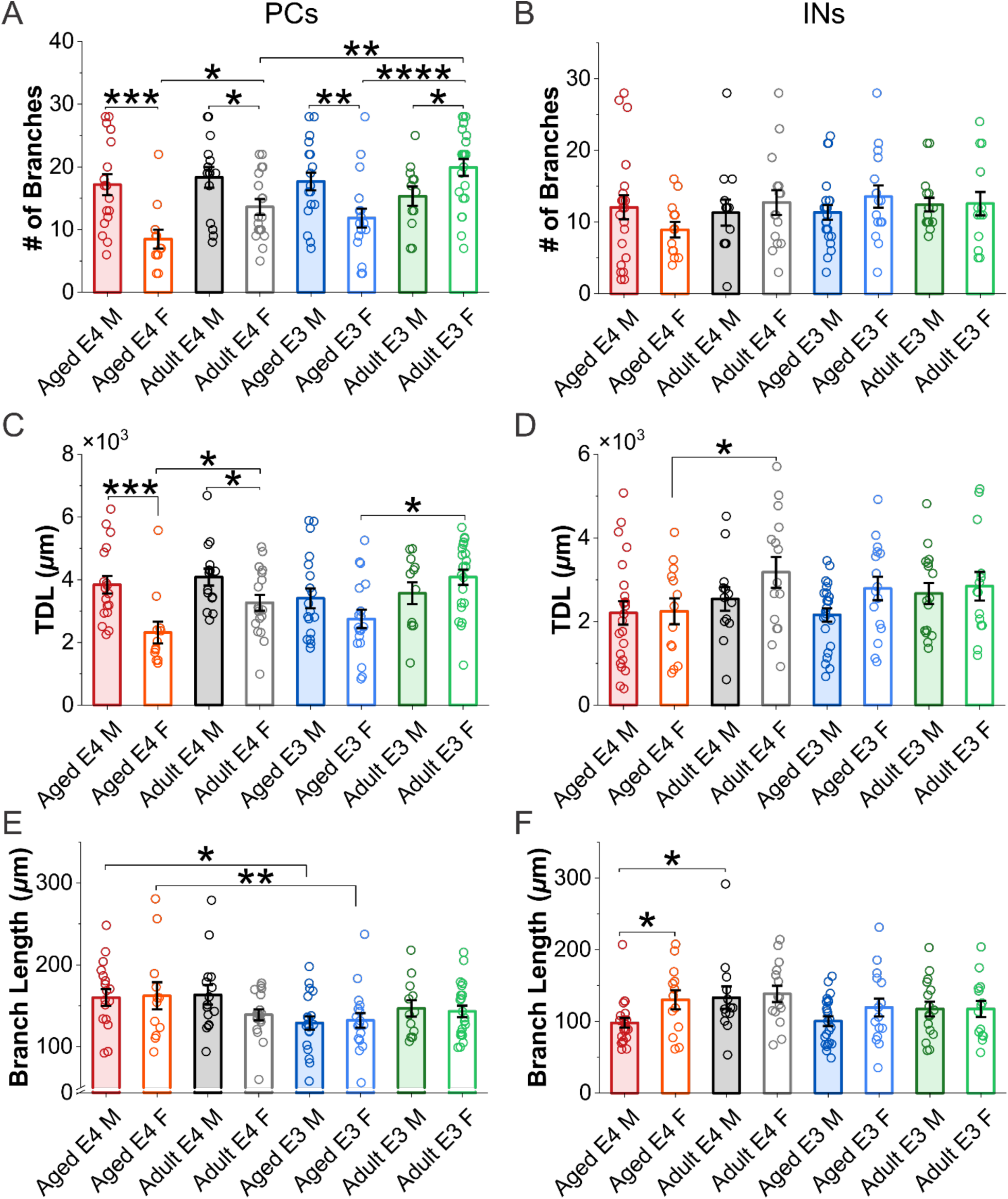
Influence of genotype, age, and sex on the dendritic architecture. **A&B**, comparison of number of dendritic branches in PCs (A) and INs (B) across groups stratified by genotype, age, and sex. **C&D**, comparison of total dendritic length in PCs (C) and INs (D) across groups. **E&F**, comparison of average branch length in PCs (E) and INs (F) across groups. Sample sizes (cells/group): aged APOE4 (E4) males (n=18 PCs & 22 INs), aged E4 females (n=12 PCs & 13 INs), adult E4 males (n=15 PCs & 13 INs), adult E4 females (n=18 PCs & 14 INs), aged APOE3 (E3) males (n=19 PCs & 23 INs), aged E3 females (n=18 PCs & 16 INs), adult E3 males (n=12 PCs & 16 INs), and adult E3 females (n=22 PCs & 14 INs). Bar graphs represent group means ± SEM with individual data points represented by symbols. Three-way ANOVA with Holm–Bonferroni correction, *p < 0.05, **p < 0.01, ***p < 0.001, ****p < 0.0001.

Because branch number reports only how often the arbor divides, we next examined TDL to determine whether the overall extent of the arbor followed the same rule. It did indeed: three-way ANOVA showed that TDL of PCs was subject to significant effects of age (F(1,126) = 10.29, p = 0.00169), sex (F(1,126) = 8.86, p = 0.0035), and age × sex (F(1,126) = 5.03, p = 0.0267) or genotype × sex (F(1,126) = 6.89, p = 0.00975) interactions (Tab. 1). Specifically, TDL declined with age in females but not males of both genotypes (2314 ± 343 µm in aged E4, n = 12 vs 3264 ± 252 µm in adult E4, n = 18, a 29.1% reduction, t = −2.21, p = 0.02862; 2748 ± 294 µm in aged E3, n = 18 vs 4081 ± 243 µm in adult E3, n = 22, a 32.7% reduction, t = −3.64, p = 0.00039; Fig. 2C). E4 males had longer TDL than E4 females in both aged (3840 ± 281 µm, n = 18 vs 2314 ± 343 µm, n = 12; t = 3.56, p = 0.00053) and adult (4085 ± 269 µm, n = 15 vs 3264 ± 252 µm, n = 18; t = 2.04, p = 0.04351) cohorts (Fig. 2C), indicating sex effects across ages. However, the TDL of INs was only influenced by age (F(1,123) = 5.90, p = 0.01658) and sex (F(1,123) = 3.97, p = 0.04853) (Tab. 1). This was supported by post hoc paired comparison showing shorter TDL in aged compared to adult E4 females (2247 ± 309 µm, n = 13 vs 3302 ± 373 µm, n = 14; t = −2.50, p = 0.01357) (Fig. 2D).

Thus, TDL is also subject to the influence from APOE genotype, age, sex and their interactions mainly in PCs.

Alterations in TDL could be just due to the variations in the number of branches in each cell or the net interactive effects between the number of branches and the average branch length. To distinguish these possibilities, we measured the average branch length in each cell and compared the average among groups stratified by APOE genotype, age and sex. The ANOVA analysis showed that branch length in PCs was impacted by APOE genotype alone (F(1,126) = 7.04, p = 0.00901; Tab. 1), with no significant effects of interactions among the other two factors. Post hoc paired comparison revealed that aged E4 mice had longer branches than aged E3 mice in both sexes (males: 160.1 ± 10.1 µm, n = 18 vs 128.9 ± 8.2 µm, n = 19, a 24.3% increase, t = 2.49, p = 0.01413; females: 162.1 ± 16.5 µm, n = 12 vs 132.1 ± 9.0 µm, n = 18, a 22.7% increase, t = 2.11, p = 0.03706; Fig. 2E). In INs, the average branch length was altered by both age (F(1,123) = 4.44, p = 0.03722) and sex (F(1,123) = 4.16, p = 0.04341) (Tab. 1). Consistently, post hoc paired comparison showed that aged E4 males had shorter branches than both adult E4 males (98.0 ± 6.7 µm, n = 22 vs 133.2 ± 15.7 µm, n = 13; t = −2.46, p = 0.01512) and aged E4 females (98.0 ± 6.7 µm vs 130.2 ± 13.3 µm, n = 13; t = −2.26, p = 0.0257) (Fig. 2F). These results demonstrated that the average branch length was the only morphological parameter exclusively affected by APOE genotype in PCs but decreased with age in E4 males in INs.

Collectively, the morphology of PCs was subject to more profound and complex influence from APOE genotype, age and sex than INs. APOE genotype exerted the most cell type–specific effects: its interaction with age shaped somatic size whereas its interaction with sex altered the number of branches selectively in PCs. Aging reduced dendritic dimensions in both cell types. Sex influenced dendritic dimensions in both cell types, and in PCs it did so in interaction with APOE genotype and age.

### Effects of APOE genotype, age, and sex on neuron membrane properties

Given the observed impact of APOE genotype, age, and sex on AON neuronal morphology that determines neuronal function, we next asked how these factors influenced membrane properties by focusing on resting membrane potential (RMP) and input resistance (R_in_) both in PCs and INs (Fig. 3).

**Figure 3.**
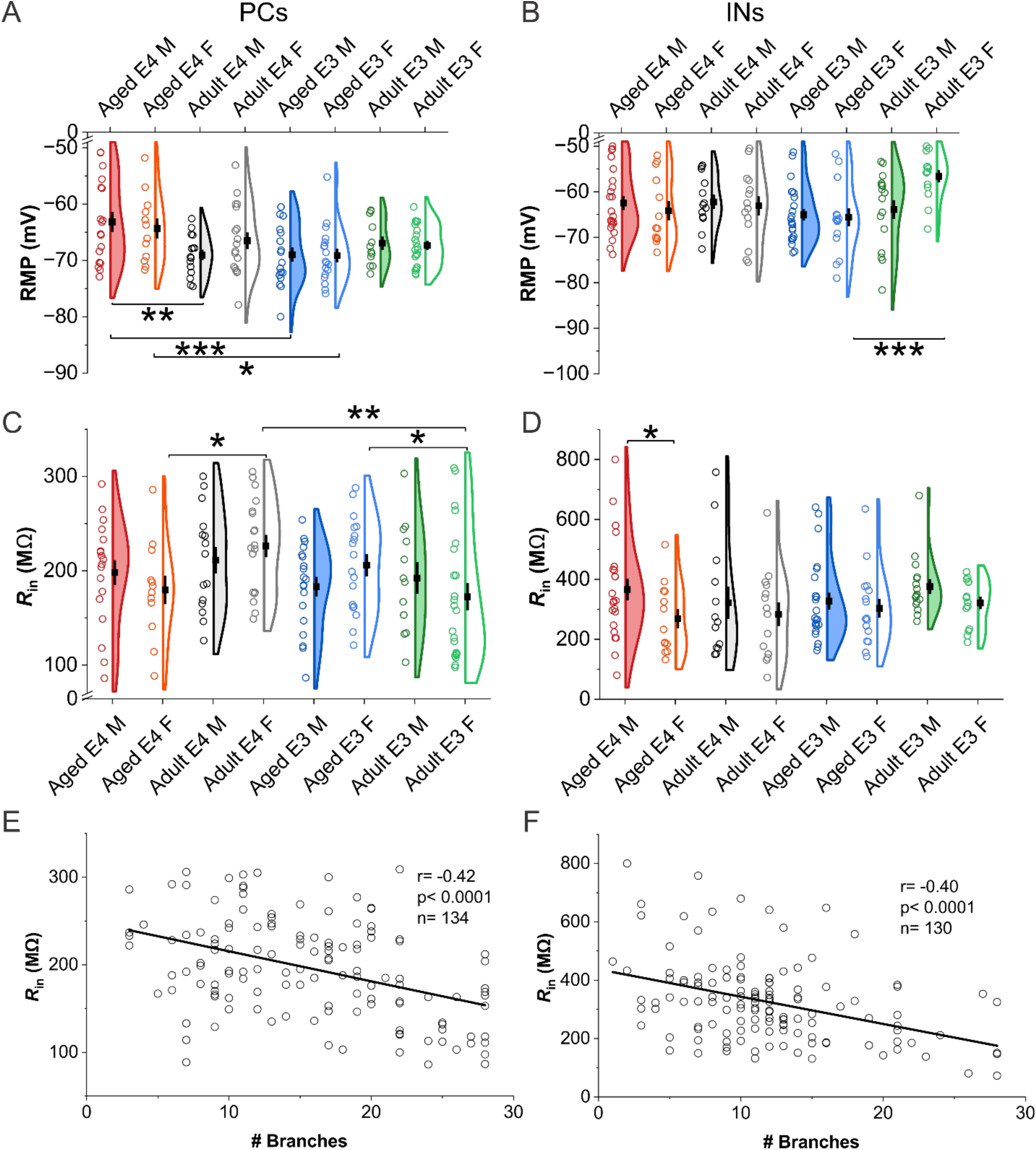
Impact of genotype, age, and sex on membrane electrophysiological properties. **A &B**, comparison of resting membrane potential (RMP) in PCs (A) and INs (B) among groups stratified by genotype, age, and sex. **C&D**, Comparison of input resistance (R_in_) in PCs (C) and INs (D) among groups. **E&F,** relationship between the input resistance and number of dendritic branches (linear fit) in PCs (E; Pearson’s r = −0.42, p < 0.0001, n = 134) and INs (F; r = −0.40, p < 0.0001, n = 131). Sample sizes (cells/group): aged APOE4 (E4) males (n=18 PCs & 22 INs), aged E4 females (n=12 PCs & 13 INs), adult E4 males (n=15 PCs & 13 INs), adult E4 females (n=18 PCs & 14 INs), aged APOE3 (E3) males (n=19 PCs & 23 INs), aged E3 females (n=18 PCs & 16 INs), adult E3 males (n=12 PCs & 16 INs), and adult E3 females (n=22 PCs & 14 INs). Violin graphs represent means ± SEM with superimposed individual data points. Statistical analysis in A-D was performed using three-way ANOVA with Holm–Bonferroni post hoc correction. *p < 0.05, **p < 0.01, ***p < 0.001.

RMP in PCs was influenced by APOE genotype (F(1,126) = 6.10, p = 0.01487) together with a significant age × genotype interaction (F(1,126) = 9.87, p = 0.0021; Tab. 1).

These effects were further supported by post hoc paired comparisons. Aged E4 mice had a more depolarized RMP than their aged E3 counterparts across sexes (males: −63.2 ± 1.7 mV, n = 18 vs −69.0 ± 1.2 mV, n = 19, a 5.8 mV difference, t = 3.41, p = 0.00088; females: −64.4 ± 1.8 mV, n = 12 vs −69.1 ± 1.2 mV, n = 18, a 4.8 mV difference, t = 2.47, p = 0.0148; Fig. 3A). But no genotype difference was detected in adult animals, suggesting age-dependence. Furthermore, RMP in PCs of aged E4 males was relatively depolarized compared to that in adult E4 males (−63.2 ± 1.7 mV, n = 18 vs −69.1 ± 0.9 mV, n = 15; t = 3.25, p = 0.0015; Fig. 3A). This age-related difference was detected only in E4 males, and not in E4 females or in E3 mice of either sex. Collectively, effects of APOE genotype on RMP in PCs were only present in aged mice, in which E4 mice showed a depolarized RMP relative to E3 mice. The post hoc pairwise comparison showed age difference in E4 males, in which the RMP in aged mice was depolarized relative to adults. Similarly, RMP in INs was also influenced by age (F(1,123) = 5.12, p = 0.02535) (Tab. 1) but only in E3 females. Specifically, aged E3 females showed more hyperpolarized RMP than adult E3 females (−65.6 ± 2.0 mV, n = 16 vs −56.6 ± 1.4 mV, n = 14, a 9.0 mV difference; t = −3.58, p = 0.0005; Fig. 3B).

In contrast to RMP, R_in_ in PCs was subject to no main effect of genotype, age, or sex (Tab. 1) but an influence from the age × genotype interaction (F(1,126) = 4.73, p = 0.03156; Tab. 1). Post hoc comparisons localized this effect to females, in which the two genotypes differed in opposite directions between the adult and aged groups: R_in_ was lower in aged than adult E4 mice (179.6 ± 15.1 MΩ, n = 12 vs 226.2 ± 11.9 MΩ, n = 18, a 20.6% reduction; t = −2.35, p = 0.02053) but higher in aged than adult E3 mice (206.0 ± 11.7 MΩ, n = 18 vs 172.5 ± 14.5 MΩ, n = 22, a 19.4% increase; t = 1.98, p = 0.04975; Fig. 3C). As a result, adult E4 females had markedly higher R_in_ than adult E3 females (226.2 ± 11.9 MΩ vs 172.5 ± 14.5 MΩ; t = 3.17, p = 0.00189; Fig. 3C), a genotype difference that was absent in age groups. R_in_ in PCs is thus sculpted by APOE genotypes in a sex-and age-dependent manner rather than being uniformly altered. In contrast, R_in_ in INs was only shaped by sex alone (F(1,123) = 4.64, p = 0.0331), with no contribution of age, genotype, or interactions (Tab. 1). Specifically, aged E4 males had higher R_in_ than aged E4 females (366.2 ± 36.4 MΩ, n = 22 vs 269.2 ± 32.3 MΩ, n = 13; t = 2.05, p = 0.04227; Fig. 3D).

Beyond these group-level differences, we further asked whether R_in_ was related to dendritic architecture at the level of individual cells. R_in_ in both PCs (Pearson’s r = −0.42, p < 0.0001, n = 134; Fig. 3E) and INs (r = −0.40, p < 0.0001, n = 131; Fig. 3F) was negatively correlated with the number of dendritic branches, indicating that cells with more complex dendritic arbors have lower input resistance.

Together, these findings showed that the three AD risk factors shape membrane properties in a cell type–specific manner. APOE genotype acted exclusively on PCs, depolarizing RMP in aged animals of both sexes and reversing the age trajectory of R_in_ in females, while contributing to neither parameter in INs. There weresignificant genotype differences in R_in_ of PCs in females. Aging depolarized RMP in E4 PCs and shaped RMP in INs of only E3 mice. Sex was the sole impactor of R_in_ in INs. R_in_ scaled inversely with dendritic branching in both cell types. The net effect is that RMP was more depolarized in aged E4 PCs than in aged E3 PCs, whereas in INs it was influenced by age without a contribution of genotype.

### Genotype-, age-, and sex-dependent variations in action potential characteristics

To further examine the potential influence of same factors on membrane electrophysiological properties, we focused on action potential characteristics by measuring the amplitude of current-evoked action potentials (APs, Fig. 4A&B) and afterhyperpolarization (AHP) in both PCs and INs.

**Figure 4.**
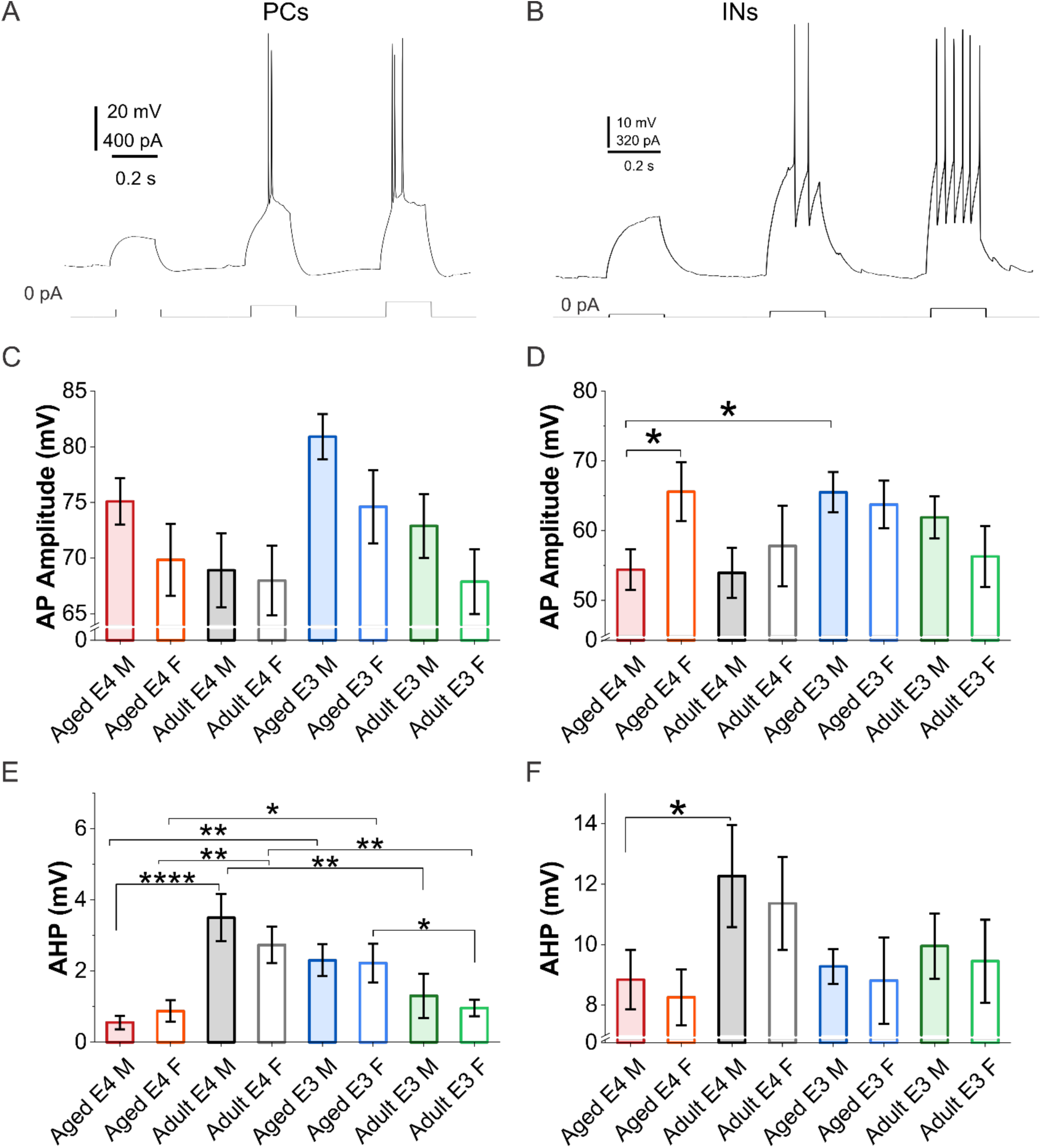
Genotype-, age-, and sex-dependent variations in action potential characteristics. **A&B**, representative current clamp recording traces (top) showing action potentials evoked by injection of progressively depolarizing currents (bottom, 300 ms/step) in a PC (A) from an adult APOE4 female mouse and in an IN (B) from an adult APOE3 male mouse. **C &D,** comparison of AP amplitude in PCs (C) and INs (D) across groups stratified by genotype, sex, and age. **E &F,** comparison of afterhyperpolarization (AHP) in PCs (E) and INs (F) across groups. Sample sizes (cells/group): aged APOE4 (E4) males (n=18 PCs & 22 INs), aged E4 females (n=12 PCs & 13 INs), adult E4 males (n=15 PCs & 13 INs), adult E4 females (n=18 PCs & 14 INs), aged APOE3 (E3) males (n=19 PCs & 23 INs), aged E3 females (n=18 PCs & 16 INs), adult E3 males (n=12 PCs & 16 INs), and adult E3 females (n=22 PCs & 14 INs). Bar graphs represent mean ± SEM of each group. Statistical comparisons were performed using three-way ANOVA with Holm–Bonferroni correction. *p < 0.05, **p < 0.01, ***p < 0.001, ****p < 0.0001.

Although the three-way ANOVA showed that AP amplitude in PCs was influenced by age (F(1,126) = 7.40, p = 0.00744) and sex (F(1,126) = 4.35, p = 0.03901), but not by APOE genotype or their interactions (Tab. 1), post hoc individual pairwise comparison revealed no statistical significance among groups stratified by APOE genotype, age and sex (Fig. 4C), with AP amplitude ranging from 67.9 ± 2.9 mV (adult E3 females, n = 22) to 80.9 ± 2.0 mV (aged E3 males, n = 19). In contrast, AP amplitude in INs showed a significant impact from sex × genotype interaction (F(1,123) = 4.40, p = 0.03791; Tab. 1). Consistently, post hoc paired comparisons revealed that aged E4 males had smaller AP amplitudes than both aged E3 males (54.4 ± 2.9 mV, n = 22 vs 65.5 ± 2.9 mV, n = 23, a 17.0% reduction; t = −2.57, p = 0.0113) and aged E4 females (54.4 ± 2.9 mV vs 65.6 ± 4.2 mV, n = 13, a 17.0% reduction; t = −2.21, p = 0.02925; Fig. 4D). These results suggest that AP amplitude was influenced by age and sex in PCs, and by the interaction between sex and genotype in INs.

In contrast to AP amplitude, AHP amplitude in PCs was affected by genotype × age interactions (F(1,126) = 29.322, p < 0.0001, Tab. 1). Specifically, APOE genotype oppositely affected AHP depending on the age regardless of sexes. For example, aged E4 mice had smaller AHPs than aged E3 mice across sexes (males: 0.55 ± 0.19 mV, n = 18 vs 2.30 ± 0.45 mV, n = 19, a 75.9% reduction, t = −2.96, p = 0.00363; females: 0.88 ± 0.30 mV, n = 12 vs 2.23 ± 0.54 mV, n = 18, a 60.7% reduction, t = −2.02, p = 0.04562) while adult E4 mice had larger AHPs than their E3 counterparts regardless of sexes (males: 3.50 ± 0.66 mV, n = 15 vs 1.30 ± 0.62 mV, n = 12, t = 3.17, p = 0.00194; females: 2.73 ± 0.51 mV, n = 18 vs 0.96 ± 0.23 mV, n = 22, t = 3.11, p = 0.00233) (Fig. 4E). On the other hand, AHP was significantly reduced in aged E4 mice compared to adult animals across sexes (males: 0.55 ± 0.19 mV vs 3.50 ± 0.66 mV, an 84.2% reduction, t = −4.70, p < 0.0001; females: 0.88 ± 0.30 mV vs 2.73 ± 0.51 mV, a 67.9% reduction, t = −2.78, p = 0.00628) but was elevated in aged E3 females relative to adult E3 females (2.23 ± 0.54 mV, n = 18 vs 0.96 ± 0.23 mV, n = 22; t = 2.22, p = 0.02847; Fig. 4E). In INs, however, AHP was influenced by age alone (F(1,123) = 5.43, p = 0.02143) (Tab. 1). This effect was confined to E4 males in which aged mice had smaller AHPs than adult ones (8.84 ± 0.98 mV, n = 22 vs 12.27 ± 1.69 mV, n = 13, a 28.0% reduction; t = −2.15, p = 0.03389; Fig. 4F).

Taken together, these results support that the three risk factors differentially regulate AP characteristics depending on neuron types. AP amplitude was influenced by age and sex in PCs, whereas in INs it was subject to influence of sex × genotype interactions, with lower amplitude in aged E4 males than in aged E4 females and aged E3 males.

AHP in PCs was differentially affected by APOE genotype and age, with genotype effects depending on age and vice versa, suggesting their interaction. In INs, however, AHP was only subject to age influence in E4 males, with attenuation in aged mice compared to adult mice.

### Variations in sEPSC frequency across genotypes, ages, and sexes

Impact on neuron intrinsic properties may lead to alterations in synaptic transmission in local circuits. To test this, we first recorded spontaneous excitatory postsynaptic currents (sEPSCs) in PCs and INs. These currents were mediated by AMPA receptors, as evidenced by their complete abolition following bath application of DNQX. (Fig. 5A).

**Figure 5.**
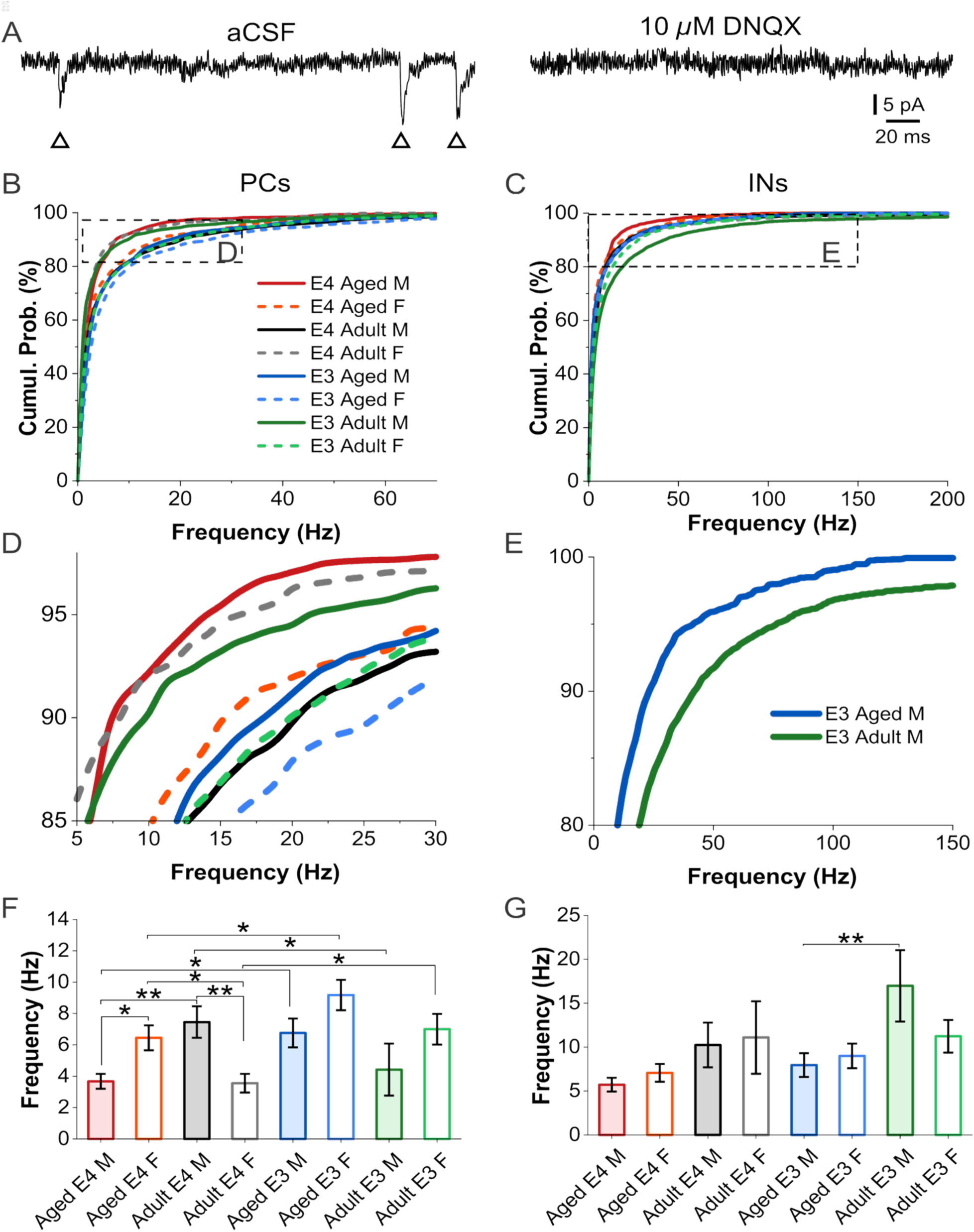
Variations in sEPSC frequency across genotypes, ages, and sexes. **A**, Representative recording traces showing spontaneous excitatory postsynaptic currents (sEPSCs, indicated by triangle symbols) recorded from a pyramidal neuron voltage clamped at −70 mV in the presence of ACSF (left) and after DNQX treatment for 10 min (right). **B &C**, comparison of cumulative probability (Cumul. Prob.) of sEPSC frequency distributions in PCs (B) and INs (C) across groups stratified by genotype, sex, and age. **D&E**, magnified view of the dashed boxed region in B & C. **F&G**, quantitative comparison of the average sEPSC frequency in PCs (F) and INs (G) across groups. Sample sizes (cells/group): aged APOE4 (E4) males (n = 13 PCs & 21 INs), aged E4 females (n = 12 PCs & 13 INs), adult E4 males (n = 15 PCs & 13 INs), adult E4 females (n = 12 PCs & 14 INs), aged APOE3 (E3) males (n = 17 PCs & 21 INs), aged E3 females (n = 14 PCs & 16 INs), adult E3 males (n = 11 PCs & 16 INs), and adult E3 females (n = 17 PCs & 14 INs). Bar graphs represent means ± SEM. Statistical analysis was performed using three-way ANOVA with Holm–Bonferroni post hoc correction. *p < 0.05, **p < 0.01.

Among aged animals, the cumulative probability distributions of sEPSC frequencies in PCs of both E4 males (solid red curve) and E4 females (dashed red curve) were shifted leftward relative to their E3 counterparts (blue solid and dashed curves), respectively (Fig 5 B&D), indicating fewer sEPSCs in aged E4 mice regardless of sex. In adult mice, although the same metric in E4 females (dashed gray curve) similarly shifted leftwards compared to E3 females (dashed green curve), it shifted rightwards in E4 males (solid black curve) relative to E3 males (solid green curve), indicating opposite genotype effects between sexes.

In line with these interpretations, the comparison of the mean sEPSC frequency in PCs among different groups stratified by APOE genotype, age and sex revealed significant influence from genotype (F(1,103) = 5.04, p = 0.02694), age × sex (F(1,103) = 5.52, p = 0.02074), sex × genotype (F(1,103) = 4.87, p = 0.02959), and genotype × age × sex (F(1,103) = 6.08, p = 0.01537) interactions (Tab. 1). Specifically, aged E4 mice showed lower sEPSC frequencies than aged E3 mice across sexes (males: 3.67 ± 0.47 Hz, n = 13 vs 6.76 ± 0.92 Hz, n = 17, a 45.7% reduction, t = −2.41, p = 0.01775; females: 6.45 ± 0.80 Hz, n = 12 vs 9.18 ± 0.97 Hz, n = 14, a 29.7% reduction, t = −2.00, p = 0.04872).

However, in adult mice, although similar APOE genotype effect was observed in females (3.56 ± 0.60 Hz, n = 12 vs 7.00 ± 0.98 Hz, n = 17, a 49.2% reduction, t = −2.63, p = 0.00995), E4 males had higher sEPSC frequency than E3 males (7.46 ± 1.01 Hz, n = 15 vs 4.42 ± 1.66 Hz, n = 11, a 68.6% increase, t = 2.20, p = 0.03019), indicating sex-dependent APOE genotype effects (Fig. 5F). Age effects were observed only in E4 mice, in which sEPSC frequency declined with age in males (3.67 ± 0.47 Hz vs 7.46 ± 1.01 Hz; t = −2.87, p = 0.00497) but rose with age in females (6.45 ± 0.80 Hz vs 3.56 ± 0.60 Hz; t = 2.04, p = 0.04416), suggesting sex-dependence. In contrast, sex differences were present in both aged and adult E4 mice but with different patterns.

Specifically, aged E4 males had a lower average sEPSC frequency than aged E4 females (3.67 ± 0.47 Hz in males vs 6.45 ± 0.80 Hz in females; t = −1.99, p = 0.04882) while adult E4 mice showed an opposite sex influence (7.46 ± 1.01 Hz in males vs 3.56 ± 0.60 Hz in females; t = 2.90, p = 0.00462), indicating age-dependence.

In contrast to PCs, 3-way ANOVA revealed merely age differences in sEPSC frequency in INs (F(1,120) = 8.78, p = 0.00367, Tab. 1). This was evidenced by a rightward shift in the cumulative probability distribution of sEPSC frequency (Fig 5C & 5E, solid green curve vs solid blue curve) and a higher mean sEPSC frequency in adult E3 males compared to aged E3 males (16.98 ± 4.07 Hz, n = 16 vs 7.95 ± 1.35 Hz, n = 21, a 53.2% reduction with age; t = −3.02, p = 0.00308) (Fig 5G), suggesting an age-associated reduction of excitatory synaptic drive in E3 males.

Overall, the sEPSC frequency was predominantly altered in PCs, where it was subject to influence of APOE genotype, age, sex, and their interactions. In INs, sEPSC frequency was influenced only by age, with reduction in aged compared to adult E3 males.

### Variations in sEPSC amplitude across genotypes, ages, and sexes

In contrast to sEPSC frequency, sEPSC amplitude in PCs was affected only by age (F(1,103) = 6.55, p = 0.01196). This is evidenced by a leftward shifted cumulative probability distribution (dashed blue curve vs dashed green curve, Fig 6A & 6C) and a smaller average amplitude of sEPSCs in aged E3 female mice compared to adult E3 female mice (8.70 ± 0.45 pA, n = 14 vs 12.13 ± 0.96 pA, n = 17, a 28.3% reduction; t =−2.52, p = 0.0132, Fig 6E).

**Figure 6.**
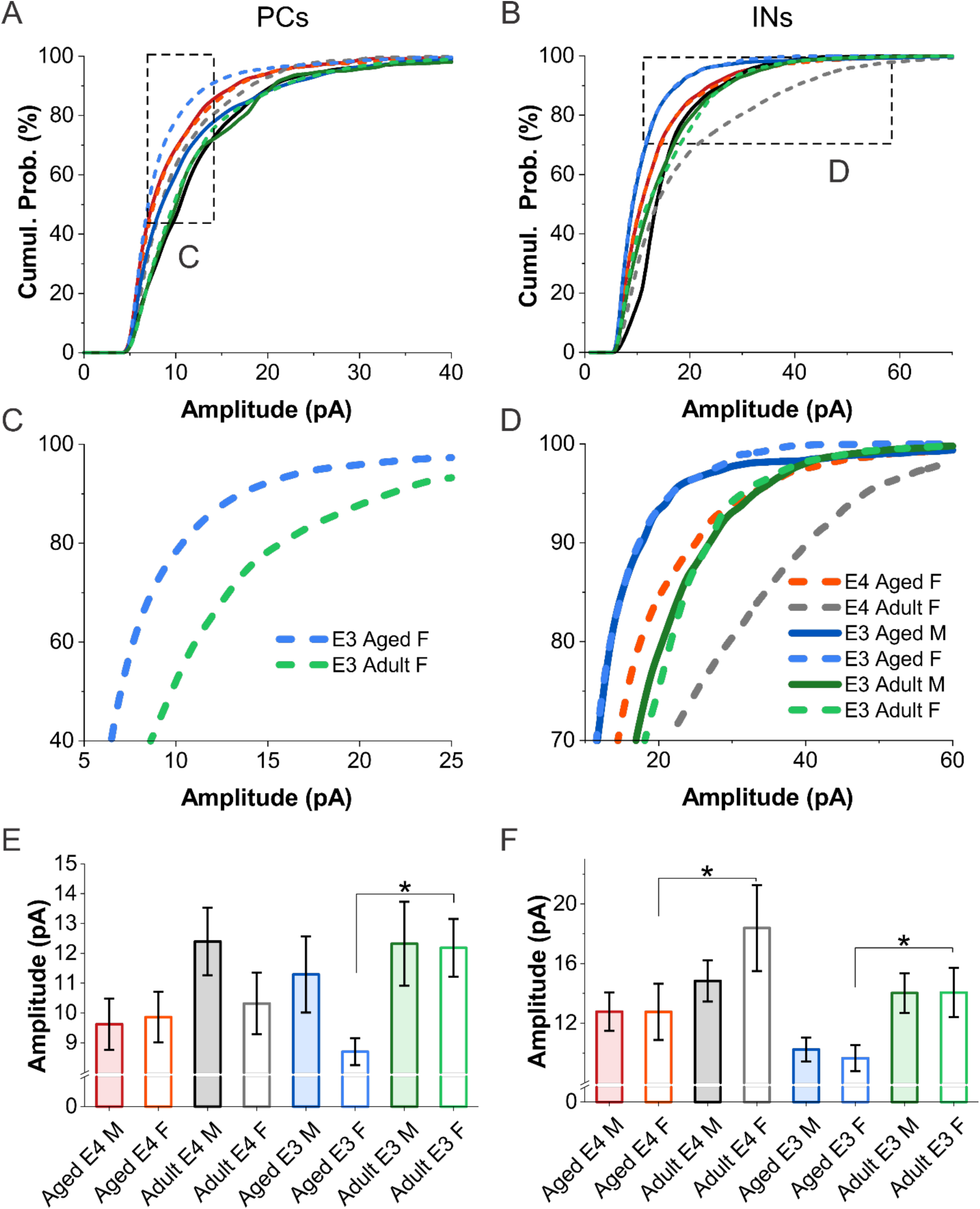
Variations in sEPSC amplitude across genotypes, ages, and sexes. **A &B**, comparison of cumulative probability (Cumul. Prob.) of sEPSC amplitude distributions in PCs (A) and INs (B) across groups stratified by genotype, sex, and age. **C&D**, magnified view of the dashed boxed region in A & B, respectively. **E&F**, quantitative comparison of the average sEPSC amplitude in PCs (E) and INs (F) across groups. Sample sizes (cells/group): aged APOE4 (E4) males (n = 13 PCs & 21 INs), aged E4 females (n = 12 PCs & 13 INs), adult E4 males (n = 15 PCs & 13 INs), adult E4 females (n = 12 PCs & 14 INs), aged APOE3 (E3) males (n = 17 PCs & 21 INs), aged E3 females (n = 14 PCs & 16 INs), adult E3 males (n = 11 PCs & 16 INs), and adult E3 females (n = 17 PCs & 14 INs). Bar graphs represent means ± SEM. Statistical analysis was performed using three-way ANOVA with Holm–Bonferroni post hoc correction. *p < 0.05, **p < 0.01.

In contrast to PCs, more differences were observed in sEPSC amplitude in INs. As shown in Fig 6B & 6D, the cumulative distributions of sEPSC amplitude were generally separated by age, with aged groups (dashed red curve, dashed and solid blue curves) shifted leftward to their adult counterparts (dashed gray curve, dashed and solid green curves), and by genotype, with E4 distributions (dashed red and gray curves) shifted to the right of E3 counterparts (dashed blue and dashed green curves). These genotype (F(1,120) = 6.08, p = 0.01507) and aging (F(1,120) = 13.18, p = 0.00042) effects were confirmed by three-way ANOVA analysis of the average sEPSC amplitude among groups (Tab. 1). Post hoc paired comparison further revealed that sEPSC amplitude declined with age in females across APOE genotypes (E4: 12.76 ± 1.88 pA, n = 13 vs 18.38 ± 2.88 pA, n = 14, a 30.6% reduction, t = −2.48, p = 0.01458; E3: 9.65 ± 0.88 pA, n = 16 vs 14.06 ± 1.65 pA, n = 14, a 31.4% reduction, t = −2.05, p = 0.04296; Fig. 6F).

These results suggest that spontaneous excitatory synaptic input in INs declines with age in females regardless of APOE genotypes.

Collectively, these findings show that APOE genotype, age, and sex shape spontaneous excitatory transmission differently in both neuron types but in different patterns. In PCs, genotype influenced frequency through a three-way interaction with age and sex, whereas amplitude was affected primarily by age. In INs, both frequency and amplitude were attenuated by aging. Thus, E4 was associated with lower sEPSC frequency in aged PCs of both sexes while both sEPSC amplitude and frequency were reduced in aged compared with adult INs across APOE genotypes.

### No effect of genotypes, ages, and sexes on sIPSC frequency

Given the observed modulation of structural and intrinsic membrane properties in interneurons by APOE genotype, age and sex, we next try to answer whether these effects can be reflected at the synaptic level. To do that, we recorded spontaneous inhibitory postsynaptic currents (sIPSCs) in PCs and INs by voltage clamping them at 0 mV, the reversal potential of AMPA receptor-mediated sEPSCs. This allowed us to minimize sEPSC contamination and enhance the GABA_A_ receptor-mediated sIPSCs as shown by bath application of the selective GABA_A_ receptor blocker gabazine, which completely abolished the spontaneous outward currents (Fig. 7A).

**Figure 7.**
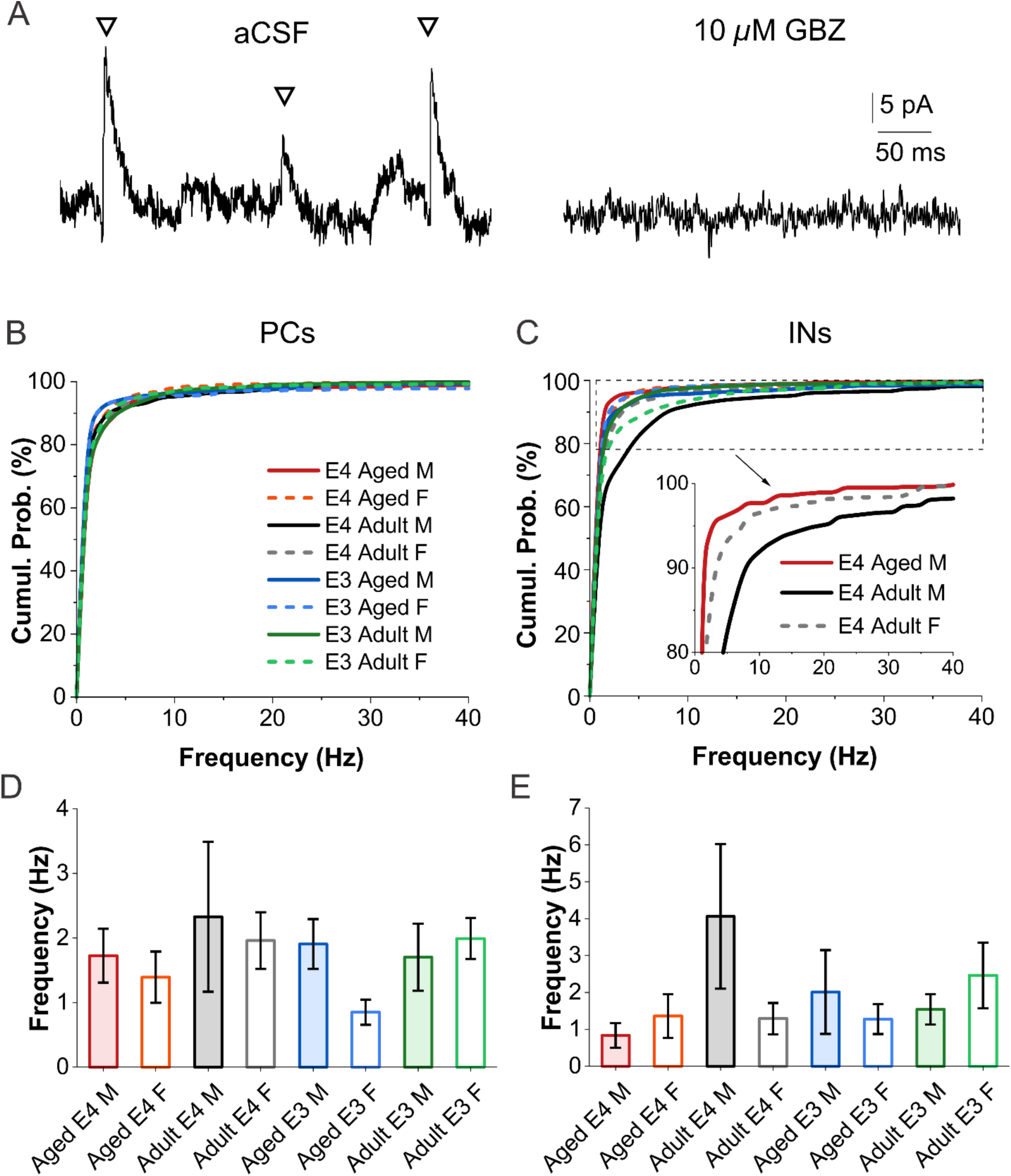
Alterations in sIPSC frequency across genotypes, ages, and sexes. **A**, Representative traces showing spontaneous inhibitory postsynaptic currents (sIPSCs, denoted by triangle symbols) recorded from a pyramidal neuron voltage clamped at 0 mV in the presence of ACSF (left) or 10 min after gabazine (GBZ) treatment (right). Note: GBZ abolished sIPSCs, confirming their mediation by GABA_A_ receptors. **B& C**, comparison of cumulative probability of sIPSC frequency in PCs (B) and INs (C) across groups stratified by genotype, sex, and age. **D & E**, quantitative comparison of sIPSC frequency in PCs (D) and INs (E) across groups. Sample sizes (cells/group): aged APOE4 (E4) males (n = 18 PCs & 15 INs), aged E4 females (n = 11 PCs & 11 INs), adult E4 males (n = 15 PCs & 10 INs), adult E4 females (n = 18 PCs & 12 INs), aged APOE3 (E3) males (n = 18 PCs & 21 INs), aged E3 females (n = 18 PCs & 15 INs), adult E3 males (n = 11 PCs & 16 INs), and adult E3 females (n = 18 PCs & 13 INs). Bar graphs represent means ± SEM. Statistical analysis was performed using three-way ANOVA with Holm–Bonferroni post hoc correction. *p < 0.05, **p < 0.01.

In PCs, the cumulative probability distributions of sIPSC frequency overlapped substantially across groups (Fig. 7B), indicating no influence from APOE genotype, age, and sex. This was confirmed by the comparison of average sIPSC frequencies among groups with three-way ANOVA (Tab. 1) and post hoc pairwise comparisons (Fig. 7D).

Similarly, although the cumulative distributions of sIPSC frequency in INs of the adult E4 males (solid black curve) was right-shifted compared to the aged E4 males (solid red curve) and adult E4 females (dashed gray curve) (Fig 7C), 3-way ANOVA showed no impact of APOE genotype, age or sex on sIPSC frequency (Tab. 1) (Fig 7E).

Collectively, sIPSC frequency in neither PCs nor INs was influenced by APOE genotype, age and sex.

### Selective influence of APOE genotypes on sIPSC amplitude

In contrast to sIPSC frequency, the cumulative distributions of sIPSC amplitude in PCs separated clearly by genotype, with the E4 distributions (red curves, Fig 8A inset) shifted rightward to the E3 distributions (blue curves, Fig 8A inset) in both sexes of aged animals (Fig. 8A). This was consistent with 3-way ANOVA showing that sIPSC amplitude was strongly influenced by genotype (F(1,119) = 14.06, p = 0.00028, Tab. 1). Specifically, aged E4 mice had larger sIPSC amplitudes than their aged E3 counterparts across sexes (males: 48.5 ± 3.5 pA, n = 18 vs 33.3 ± 1.6 pA, n = 18, a 45.8% increase, t = 3.09, p = 0.00249; females: 45.9 ± 6.2 pA, n = 11 vs 33.0 ± 2.1 pA, n = 18, a 39.3% increase, t = 2.29, p = 0.02366; Fig. 8C). In INs, although 3-way ANOVA showed that sIPSC amplitude was influenced by APOE genotype (F(1,105) = 5.01, p = 0.02727; Tab. 1), no individual pairwise comparison reached significance (Fig 8B & 8D).

**Figure 8.**
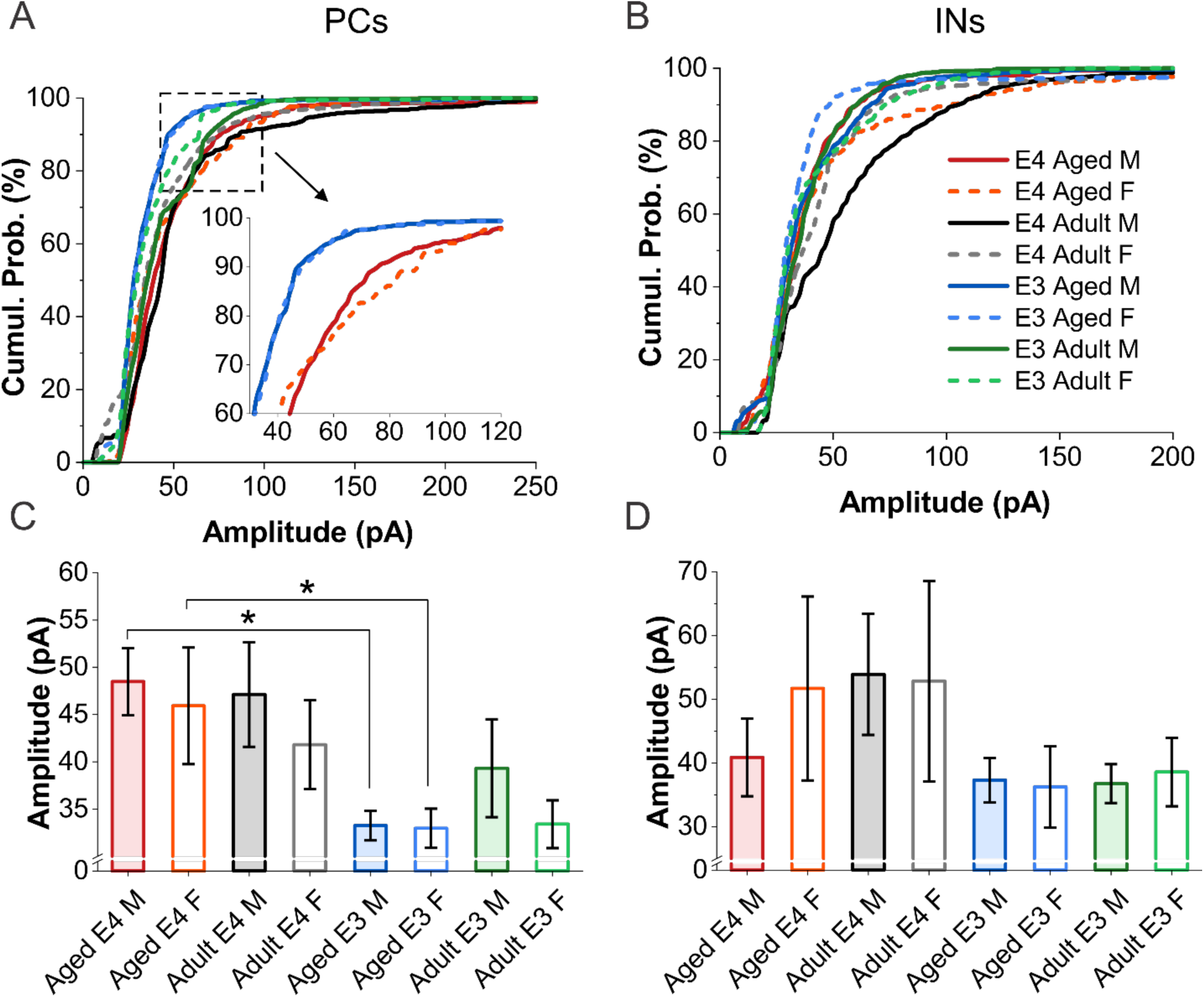
Variations in sIPSC amplitude across genotypes, ages, and sexes. **A&B**, comparison of cumulative probability of sIPSC amplitude in PCs (A) and INs (B) across groups stratified by genotype, sex, and age. **C&D**, quantitative comparison of sIPSC amplitude in PCs (C) and INs (D) across groups. Sample sizes (cells/group): aged APOE4 (E4) males (n = 18 PCs & 15 INs), aged E4 females (n = 11 PCs & 11 INs), adult E4 males (n = 15 PCs & 10 INs), adult E4 females (n = 18 PCs & 12 INs), aged APOE3 (E3) males (n = 18 PCs & 21 INs), aged E3 females (n = 18 PCs & 15 INs), adult E3 males (n = 11 PCs & 16 INs), and adult E3 females (n = 18 PCs & 13 INs). Bar graphs represent means ± SEM. Statistical analysis was performed using three-way ANOVA with Holm–Bonferroni post hoc correction. *p < 0.05, **p < 0.01.

Taken together, these results demonstrated that sIPSCs were subject to modulation by APOE genotype only in aged PCs, where sIPSC amplitude was larger in E4 than E3 mice, indicating stronger synaptic inhibition.

## Discussion

To determine how APOE genotype interacts with age and sex to shape AON neuronal structure and function, we combined whole-cell recordings from PCs and INs with post hoc morphological reconstruction in humanized APOE3 and APOE4 knock-in mice.

Three findings emerged. First, E4 produced no uniform shift; effects were distributed across structural, intrinsic, and synaptic measures and sometimes occurred in opposite directions. Second, these effects differed between PCs and INs, with broader genotype-dependent remodeling in PCs than INs. Third, age and sex substantially modified the E4 phenotype, including a genotype × age × sex interaction for PC sEPSC frequency.

Thus, APOE4 produces context-dependent neuronal remodeling in the AON rather than a single gain-or loss-of-function state.

### APOE4 selectively sculpts PC structure and intrinsic excitability

Brunjes and colleagues characterized AON neurons morphologically and physiologically in rats and mice, defining pyramidal cells and several distinct interneuron classes(Brunjes and Kenerson, 2010; Brunjes et al., 2011; Kay and Brunjes, 2014). Our data match that division: PCs and INs differed in both dendritic architecture and intrinsic membrane properties in E3 animals, and the two classes also diverged in how they responded to genotype, age and sex. Those studies resolved finer subclasses and found aging effects that were not uniform across structural measures(Brunjes et al., 2011), whereas we compared broad PC and IN populations across genotype, age and sex; individual parameters are therefore not directly comparable, given differences in species, age range and background. Consistent with reports of age-and E4-associated changes in PC morphology in somatosensory cortex and hippocampal CA1(Dumanis et al., 2009; Thorwald et al., 2025), we found genotype-dependent structural remodeling in AON PCs, whereas structural changes in INs were associated primarily with age and sex. Unlike cortical and hippocampal PCs(Dumanis et al., 2009; Thorwald et al., 2025), AON PCs in E4 mice did not consistently exhibit reduced dendritic complexity or arbor extent; instead, branch length increased in aged E4 animals despite age-and sex-dependent effects on soma size. Aging also reduced TDL and branch number in PCs and TDL and branch length in INs.

Intrinsic membrane properties were likewise affected more broadly in PCs than INs. Genotype influenced PC RMP and, through its interaction with age, AHP and R_in_, but not AP amplitude; in INs, AP amplitude showed a sex × genotype interaction. Aged E4 PCs exhibited depolarized RMP and reduced AHP, changes expected to increase neuronal responsiveness. However, alterations in these intrinsic properties cannot predict firing behavior because spike threshold, input resistance, and sodium channel availability also contribute(Platkiewicz and Brette, 2010). APOE4 effects on intrinsic excitability therefore depend strongly on neuronal identity, age, and sex.

Collectively, the present study demonstrated cell type-specific effects of APOE genotype, age, and sex on structure and intrinsic membrane properties particularly in PCs. We cannot exclude nuanced influences of these factors on interneuron subtypes (Kay and Brunjes, 2014), which warrants future effort.

### Synaptic alterations bridge cellular and network dysfunction

A recent in vivo study reported age-and sex-dependent E4 effects on AON firing and network dynamics in the same humanized APOE lines(Uzun et al., 2025). Because AON activity in awake mice is shaped by long-range inputs and local circuitry(Brunjes et al., 2005; Brunert et al., 2023), these changes likely reflect synaptic contributions, particularly from extrinsic sources. Because PCs substantially outnumber INs(Brunjes et al., 2005), in vivo recordings may predominantly reflect PC activity. Consistent with this, our ex vivo data show prominent intrinsic and synaptic changes in PCs. E4-associated reductions in sEPSC frequency and increases in sIPSC amplitude in aged PCs were consistent with the in vivo phenotype(Uzun et al., 2025), suggesting a shift in synaptic drive toward inhibition that could reduce spontaneous firing. The larger sIPSC amplitude may also indicate enhanced local inhibitory feedback, potentially contributing to the elevated gamma oscillations in awake E4 mice(Uzun et al., 2025), given the role of GABAergic interneurons in gamma(Bartos et al., 2007).

Complex genotype-and sex-dependent modulation of sEPSC frequency was concentrated in PCs, whereas IN sEPSC frequency was primarily affected by age. These patterns did not parallel the age-related increase in gamma oscillations in vivo, suggesting that long-range inputs, which may modulate AON interneurons(Brunjes et al., 2005; Brunert et al., 2023), also contribute to this network phenotype. The link between structural and synaptic changes remains correlative: reduced dendritic arborization with age was accompanied by reduced excitatory input, but spine density and synapse number were not measured, so no causal relationship can be drawn.

Likewise, the increased sIPSC amplitude in aged E4 PCs did not mirror the age-related pattern of sEPSC frequency and therefore cannot be interpreted simply as compensation for reduced excitation.

The E4-associated reduction in synaptic excitation and elevation in synaptic inhibition may counteract the responsiveness-favoring intrinsic changes such as depolarized RMP and reduced AHP. Thus, the net effect of APOE4 on PC output cannot be determined from the present measurements. Comparison between the ex vivo phenotype and in vivo activity also requires caution, as acute slices truncate the long-range connections that modulate AON activity in awake animals. APOE4-associated network dysfunction may therefore emerge from the integration of opposing intrinsic and synaptic alterations within a circuit shaped by long-range inputs.

### Age and sex shape the APOE4 phenotype

Age was the most pervasive modifier of neuronal properties, whereas sex frequently altered the direction of genotype effects. The clearest example was PC sEPSC frequency, where the E4 effect differed in direction between aged males and females. Sex also interacted with genotype for PC dendritic complexity and arbor extent. These findings complement in vivo evidence that females show greater AON excitability than males during adulthood, a difference that is diminished with age(Uzun et al., 2025), and epidemiological evidence for sex-dependent APOE risk(Farrer et al., 1997; Holland et al., 2013; Altmann et al., 2014; Riedel et al., 2016; Neu et al., 2017).

The cellular mechanisms underlying these sex differences remain unresolved. Sex hormones can regulate excitatory and inhibitory transmission and change with age, but hormonal status and estrous stage were not assessed, so hormonal mechanisms remain hypothetical. The cross-sectional design likewise limits interpretation of age effects as trajectories within individual animals.

The distinction between PCs and INs further shows that APOE4 does not remodel the AON uniformly. Genotype effects in PCs spanned morphology, intrinsic properties, and synaptic transmission, whereas in INs the main effects were largely those of age, influencing dendritic extent, RMP, AHP, and sEPSC frequency and amplitude. Several structural and intrinsic IN properties also changed with age and sex, and AP amplitude showed a sex × genotype interaction. Notably, aged E4 males showed convergent changes in interneuron morphology and intrinsic properties, whereas aged E4 females showed stronger genotype-dependent changes in excitatory input to PCs. Age and sex therefore determine not only the magnitude but also the cellular expression of the E4 phenotype.

### Implications for Alzheimer’s disease

The AON is affected early in Alzheimer’s disease, with tau pathology and degeneration preceding widespread neocortical involvement(Franks et al., 2015; Murray et al., 2020; Ubeda-Banon et al., 2020). APOE4-associated remodeling was detectable here in the absence of amyloid plaques and neurofibrillary tangles (data not shown). Rather than the convergent hyperexcitability described in some amyloid-based AD models(Busche and Konnerth, 2015; Zott et al., 2019), E4 altered multiple functional domains in different and sometimes opposing directions.

APOE4 may therefore establish an altered neuronal state before overt AD pathology that could influence how the AON responds to subsequent disease-related stress.

However, the present data do not establish that these changes are prodromal mechanisms of AD because amyloid and tau pathology were not introduced, and olfactory or cognitive function was not assessed. These phenotypes could represent an early substrate of vulnerability, an adaptive response to altered apoE4 function, or an APOE4-associated state unrelated to subsequent pathology. Differentiation of these possibilities and their behavioral outcomes will be interesting topics for future studies.

## Conclusions

APOE4 reshapes AON neuronal function through selective interactions with neuronal identity, age, and sex. In PCs, genotype affected dendritic structure, intrinsic excitability, and excitatory and inhibitory transmission, whereas in INs genotype main effects were confined to excitatory synaptic events. Importantly, these changes did not converge on a uniform increase or decrease in excitability: intrinsic properties in aged E4 PCs favored greater responsiveness, while synaptic changes favored reduced excitation and increased inhibition. Age and sex further determined when and in which neuronal populations these effects emerged. APOE4 therefore appears to establish a context-dependent cellular state in the AON, a potential substrate through which this early-affected circuit may differ in its response to aging and AD-related stress.

## Acknowledgments

This work was supported by R01AG069196, R01AG074216 & R01AG077541. We used an AI large language model (Claude, Anthropic) to assist with language editing of the manuscript text. We reviewed and took full responsibility for the manuscript content.

## Author Contributions

M.H. and S.L. designed research; M.H., AK and Y.L. performed research; M.H., A.K., S.B., and D.Z. analyzed data; M.H. wrote the first draft of the paper; M.H. and S.L. edited the paper; M.H. and S.L. wrote the paper.

